# Simultaneous multi-area recordings in macaque frontoparietal cortex during a manual object-interaction task

**DOI:** 10.64898/2026.09.29.754877

**Authors:** Kai Liu, Shuran Guan, Sizheng Luo, Yahui Li, Yizhi Dong, Weitao Chen, Mengmeng Chen, Dongling Zhang, Long Chen, Shundong Huang, Yulei Chen, Hongxiong Jiang, Lei Wu, Ruoyu Li, Fengmei Zhu, Yang Xie, Yanan Sui, Bin Min, Shouliang Guan, Chengyu T. Li, Tianwei Wang, Tianmeng He

## Abstract

Dexterous hand control is essential for daily activities and a top restoration priority for people with tetraplegia, yet neurophysiological datasets suitable for studying and decoding naturalistic hand behavior remain scarce. Most available recordings constrain movements to a small number of degrees of freedom or cover only a single stereotypical grasp-and-hold structure. Moreover, previous brain-computer interface (BCI) studies have largely relied on signals from a single cortical area, even though dexterous hand control is distributed across an interconnected frontoparietal network. To address these limitations, we designed a paradigm that elicits rich hand movements across 40 distinct hand-object interaction conditions. We recorded simultaneous frontoparietal activity with hand kinematics from a rhesus macaque performing them. Neural activity was recorded from the primary motor cortex, anterior intraparietal area, and ventral premotor cortex using microelectrode arrays spanning 1,920 channels, while a multi-camera markerless motion-capture system acquired video from which kinematics were extracted offline. The dataset comprises data from 20 sessions, including processed neural features, synchronized 3D coordinates of hand and elbow keypoints, and trial-level metadata including task conditions and event timing. We further provide technical validation, including a neural decoding benchmark with multiple algorithms for continuous keypoint coordination reconstruction. We found that cross-day and cross-task decoding performance varied among algorithms. Variation in channel availability leads to different performance degradation curves in different algorithms. This task-rich dataset offers a valuable resource for investigating the neural mechanisms of dexterous hand control and for advancing the performance of motor BCIs.

## Background & Summary

We publish a dataset of simultaneous frontoparietal cortical recordings and three-dimensional (3D) hand kinematics collected over 20 recording sessions from a rhesus macaque performing a manual object-interaction task. Sixteen flexible microelectrode arrays (128 channels each, 2,048 channels in total) were chronically implanted across the primary motor cortex (M1), the anterior intraparietal area (AIP), and the ventral premotor cortex (PMv). While the monkey interacted with 40 modules spanning five action categories (pull-and-push, push, button-pressing, pick-and-place, and pinch), hand-and-elbow kinematics were captured with a markerless motion-capture system. We provide (a) processed neural features (threshold crossings and spike-band power), (b) frame-synchronized 3D coordinates of 24 hand and elbow keypoints together with derived kinematic features, (c) trial-level metadata including task conditions and event timing. These data are packed in a decoding-ready, session-organized release aligning the neural features, kinematics, and trial information. With the dataset, we conducted a neural decoding benchmark for continuous keypoint coordination reconstruction with multiple algorithms.

Dexterous hand control is essential for daily activities and a high restoration priority for people with tetraplegia.^1^ Yet datasets suitable for studying and decoding naturalistic hand behavior remain scarce, because natural hand movements are remarkably complex. The anthropomorphic hand has more than 26 degrees of freedom controlled by dozens of muscles.^2,3^ Hand configurations vary systematically with object geometry^4^ and task goals.^5^ Most available neurophysiological datasets of hand control sample only a narrow slice of this complexity, either restricting movements to a few controlled degrees of freedom^6,7^ or covering many objects but only in a uniform grasp-and-hold action structure.^2^ Moreover, uniform behavioral tasks tend to yield low-dimensional manifolds of motor cortical activity, largely because such paradigms engage only a limited repertoire of motor subskills.^8^ Paradigms that enrich the temporal dynamics of behavior are therefore essential for revealing the full dimensionality of cortical population activity. By combining 40 object-interaction modules that vary in shape, size, orientation, and task demands, the present dataset extends behavioral coverage across both object and action dimensions and engages a broad repertoire of motor subskills. It enables the development and evaluation of decoding and analysis methods under conditions closer to natural hand use.

The neural substrate of dexterous control is distributed across an interconnected frontoparietal network.^3,9^ Within this network, posterior parietal areas such as AIP predominantly represent visual object properties relevant for grasping.^10^ Premotor cortex is closely linked to grip selection and action planning.^11,12^ M1 population activity covaries with the time-varying kinematics of the arm, wrist, and hand.^13^ Representations in these brain areas dynamically shift across task stages.^14^ As a result, brain-computer interfaces (BCIs) relying exclusively on single-area signals risk hitting a performance ceiling due to the omission of distributed, higher-order information.^15^ Simultaneous recordings across distributed cortical areas are valuable as they can capture a broader range of neural information across cortical areas and task stages, providing a basis for investigating how cortical networks coordinate dexterous hand movements.

Beyond neuroscientific questions, the dataset is intended to support method development. The synchronized neural and kinematic streams enable direct comparison and validation of decoding algorithms on realistic, high-dimensional data. Such resources address a long-standing need to overcome the challenges in hand movement decoding in the BCI field: clinical systems have progressed from reaching to multi-dimensional hand control but remain limited in dimensionality and accuracy.^16–19^ The accompanying markerless motion-capture pipeline and its quantified reconstruction accuracy further make the dataset a testbed for video-based behavioral analysis.

In the current study, we obtained a dataset comprising multiregional intracortical electrophysiological recordings and hand kinematics during rich hand motor tasks to support studies on dexterous hand control and motor BCIs (Figure 1A). A macaque performed 40 types of object-interaction tasks that were organized in a hierarchical structure (Figure 1B). Through the task procedure, we obtained 20 sessions of synchronized cortical recordings in M1, PMv and AIP, as well as 3D hand kinematics, together with processed features, trial metadata, and a decoding benchmark (Figure 1C). We also examined cross-day and cross-task decoding performance, as well as the robustness of decoders to reduced channel availability. The task-rich and high-throughput nature of this dataset enables examination of generalization and robustness of decoders. This dataset provides a unique resource for investigating the network-level neural mechanisms of dexterous manual control and for advancing motor BCIs.

**Figure 1.**
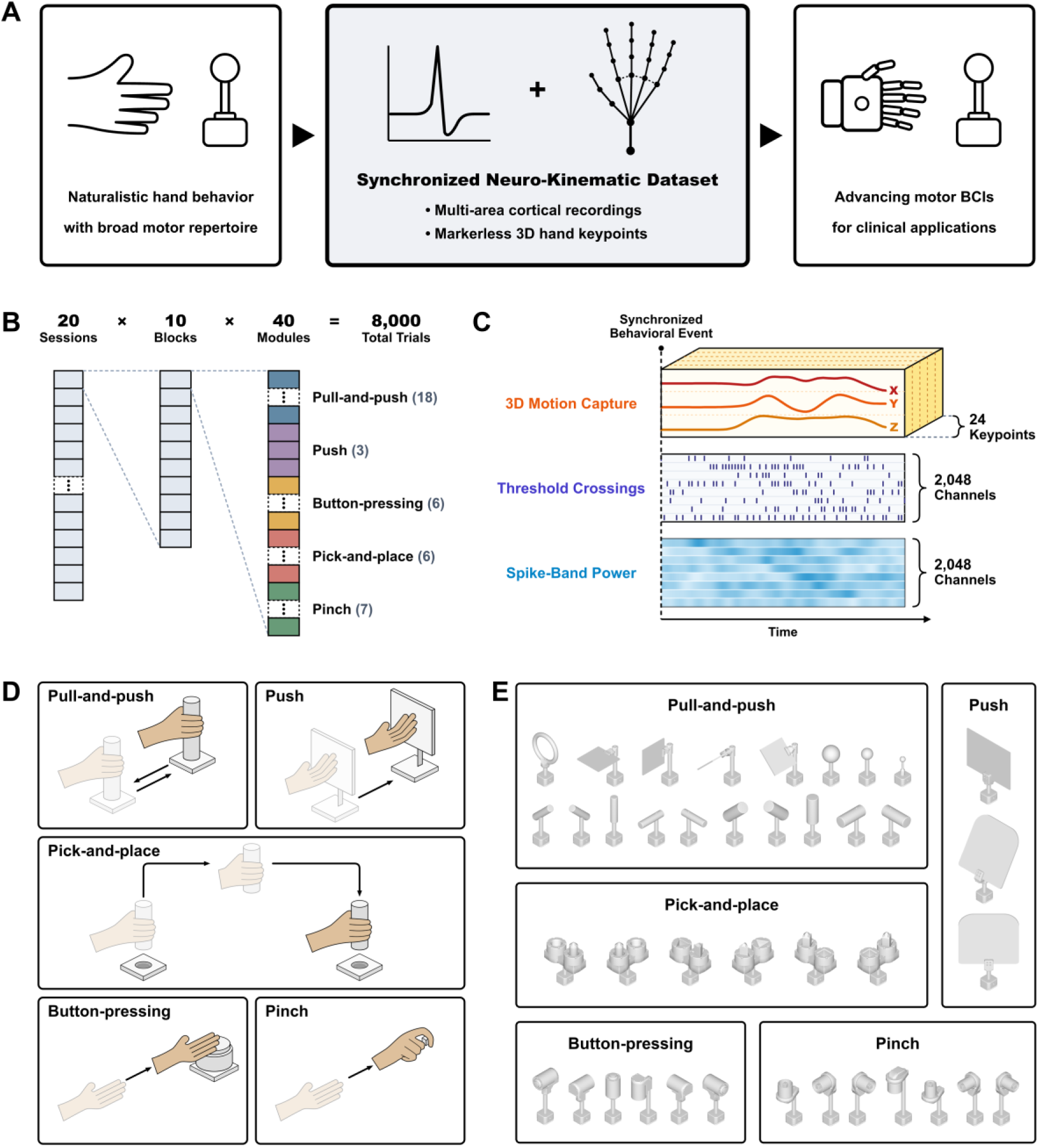
Dexterous hand movement dataset. (A) The current dataset was collected while a monkey was performing rich hand behaviors induced by multiple task conditions. It consists of synchronized intracortical electrophysiological recordings and hand kinematics. The dataset provides opportunities for studying dexterous manual control and for advancing motor BCI systems. (B) The dataset consists of 20 sessions. Each session is composed of 10 blocks. Within a block, each of the 40 modules is presented exactly once, in a pseudorandom order, yielding 40 trials per block. Thus, each session comprises 400 trials and the entire dataset comprises 8,000 trials. (C) Hand movements were captured with a motion capture system, from which 3D coordinates of 24 keypoints on the hand and forearm were extracted frame by frame. Synchronized threshold-crossing events and spike-band power were extracted from the neural recordings. (D) The behavioral paradigm consists of five task categories: pull-and-push, push, pick-and-place, button pressing, and pinch. (E) A module is a single object-configuration variant of a task. Each task category comprises multiple modules that differ in shape, size, and orientation.

## Methods

### 1. Surgical Procedures

All animal procedures were approved by the Institutional Animal Care and Use Committee of Lingang Laboratory (LGLSP-2024-024) and performed in accordance with relevant ethical guidelines. Before surgery, Monkey Q (male rhesus macaque, 8 kg) underwent 3T cranial magnetic resonance imaging (MRI) and computed tomography (CT) of the head and bilateral upper limbs. The cranial MRI provided anatomical information for surgical planning, whereas the head CT was used to design a subject-specific head post conforming to the shape of the skull. The upper-limb CT data were subsequently used to derive subject-specific limb and digit segment lengths for refining keypoint 3D reconstruction results. Two surgical procedures were then performed.

In the first surgery, a custom-designed titanium head post was affixed to the skull. It served to secure the electrode assemblies and provide stable head fixation during experimental sessions. The monkey was allowed to recover for at least three weeks after this procedure before undergoing electrode implantation.

In the second surgery, a total of 16 flexible microelectrode arrays were implanted into the primary motor cortex (M1, n = 8), anterior intraparietal area (AIP, n = 4), and ventral premotor cortex (PMv, n = 4) of the left hemisphere (Figure 2A). Each flexible microelectrode array consisted of 128 recording channels arranged in a single longitudinal column with an inter-channel spacing of 50 μm (Figure 2B). Following craniotomy and durotomy, each flexible microelectrode array was inserted into the cortex guided by a tungsten needle, reaching a depth of approximately 4-10 mm. The tungsten needle was then withdrawn. Once all electrodes had been placed, the implantation site was sealed with biocompatible silicone sealant. The last array implanted into M1 dislodged from the cortical tissue during the surgery but was still retained for signal recording. Therefore, the recorded dataset contained 2,048 channels in total, of which 1,920 channels were functional and used for subsequent analyses. Behavioral experiments began after a two-week recovery period following the second surgery.

**Figure 2.**
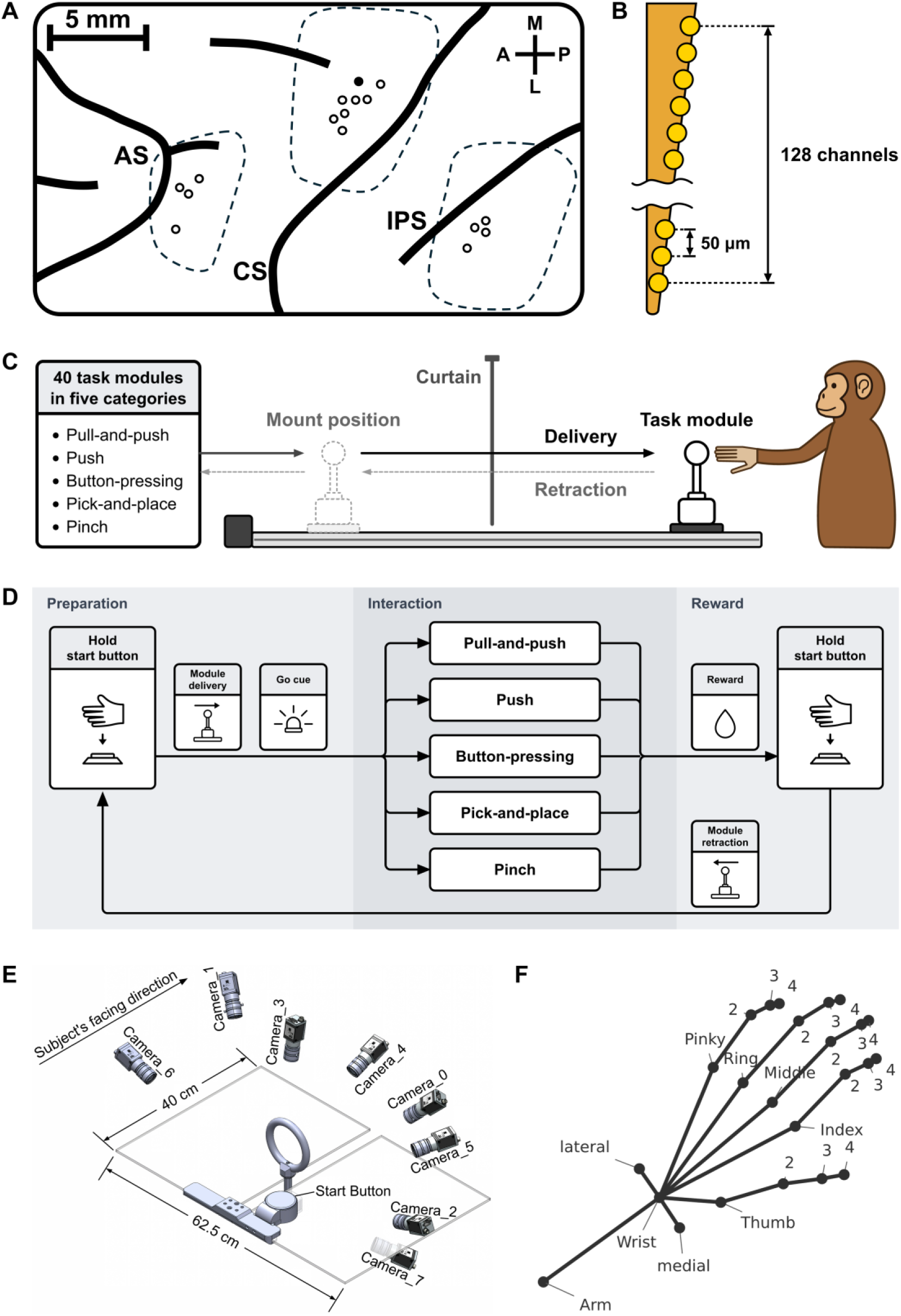
Electrode implantation, experimental setup, and task procedures. (A) Cortical microelectrode array implantation sites in Monkey Q. A total of 16 flexible microelectrode arrays (128 channels each, 2,048 channels in total) were implanted into the primary motor cortex (M1, n = 8), ventral premotor cortex (PMv, n = 4), and anterior intraparietal area (AIP, n = 4) of the left hemisphere. Dashed lines indicate the boundaries of the cranial windows. Open circles indicate functional array sites (n = 15, 1,920 channels), and the filled circle indicates the M1 array that was dislodged during surgery. Major sulcal landmarks: arcuate sulcus (AS), central sulcus (CS), and intraparietal sulcus (IPS). Scale bar, 5 mm. (B) Schematic diagram of the flexible microelectrode array. (C) Schematic diagram of the behavioral setup. The setup consists of a mounting area and the monkey’s workspace, visually isolated by a curtain and connected with a motorized rail. Task modules are mounted in the mounting area, delivered to the workspace along the rail, and retracted after the end of each trial. (D) Flowchart of the behavioral paradigm. (1) During the trial preparation stage, an experimenter mounts a pseudorandomly selected task module prompted by the control program. The monkey initiates module delivery by pressing and holding the start button until the module arrives at the workspace and a green LED illuminates as a go cue. (2) During the object interaction stage, the monkey releases the start button, reaches toward the module, and executes the manual action. (3) During the reward and end-of-trial stage, successful action completion triggers liquid reward delivery, followed by start button re-pressing and module retraction to the mounting area. (E) Approximate arrangement of the eight cameras in the motion-capture system around the operating table. The schematic is not to scale. (F) Anatomical positions and names of the 24 annotated hand and elbow keypoints for motion capture.

All surgical procedures were performed under aseptic conditions and general anesthesia (induced with Zoletil, 2-5 mg/kg, and maintained with 1.5-2% isoflurane via tracheal intubation). Throughout each surgical procedure, physiological parameters—including body temperature, heart rate, respiration rate, blood oxygen saturation, tidal volume, and end-tidal carbon dioxide—were continuously monitored. During the first week after surgery, cephalosporin antibiotics and dexamethasone were administered daily, followed by dexamethasone three times per week for long-term maintenance.

### 2. Behavioral Setup

To induce rich hand movement, we trained the monkey to interact with different modules. In each trial, the monkey was required to complete a specific interaction based on the module delivered to it. The behavioral setup consisted of a preparation area and the monkey’s workspace, visually isolated by a curtain to reduce the influence of the preparation process on the subject (Figure 2C). A one-meter-long motorized rail connected these two areas, carrying an adaptor platform which served as a mount for the task module. In each trial, a task module was mounted onto the adaptor platform in the preparation area and delivered to the workspace by the motorized rail.

A total of 40 modules across five task categories were used (Table 1, Figure 1D-E), each corresponding to a group of hand movement tasks: (1) pull-and-push tasks, where the monkey pulled the module towards itself and pushed it back, with modules differing in shape, size, and orientation; (2) push tasks, where the monkey pushed a large plate forward, with modules differing in plate orientation; (3) button-pressing tasks, where the monkey pressed a button, with modules differing in button orientation; (4) pick-and-place tasks, where the monkey retrieved a block from one slot and inserted it into another slot, with modules differing in block shape and slot arrangement; and (5) pinch tasks, where the monkey pinched a small cube and pulled it out of a transparent acrylic tube, with modules differing in tube diameter and orientation.

**Table 1.**
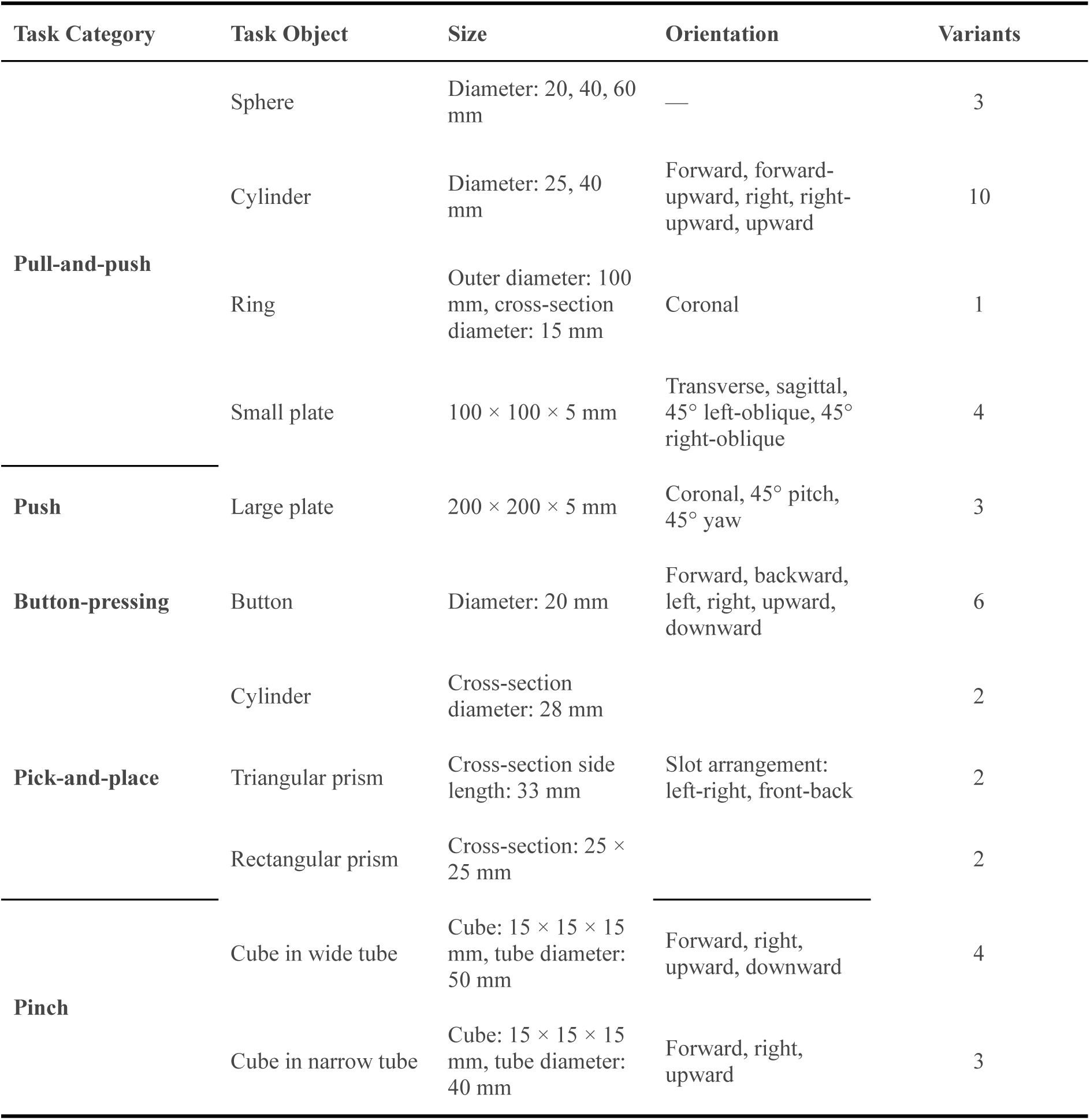
Overview of behavioral task categories and module variants.

### 3. Behavioral training and task paradigm

Before data collection, the monkey completed a seven-stage shaping protocol. In the first stage, the monkey was habituated to the experimental environment and the primate chair. Habituation was considered complete when the monkey could sit in the chair without overt signs of stress. The subsequent stages comprised training to drink from the water-delivery system, execute the pulling and pushing actions required for the pull-and-push and push modules, press buttons, pinch and extract cubes, pick up and place objects, and operate the start button to work consecutively through a full session of all the tasks. A clicker serving as a conditioned reinforcer, together with water and fruit rewards, was initially used to shape each action. Once accuracy for a trained action reached 95%, the clicker was gradually withdrawn, leaving water as the reward. Training for the seventh stage began only after accuracy across all task-module categories remained stable at or above 95%. During this stage, the monkey learned the full procedure used in the recording sessions, as detailed below.

During experimental sessions, the monkey sat in a custom primate chair with its head fixed. An opening at the front right of the chair allowed only the right hand to reach the task modules. The task proceeded through three stages: preparation, interaction and reward (Figure 2D). Before each trial, an experimenter mounted the task module specified by the control software onto the adaptor platform in the mounting area. Throughout this mounting procedure, the curtain prevented the monkey from seeing the selected module. Once the experimenter finished mounting, an auditory beep signaled the start of the trial. During the preparation stage, the monkey was required to keep its right hand on the start button near its chest. While the button was held down, the motorized rail moved the module into the workspace. The monkey maintained the start button press throughout module delivery and for an additional 100 ms after the module reached the target position to standardize the hand’s position at movement onset. The indicator LED then turned green, providing the go cue. If the monkey released the start button before the go cue, the trial was aborted immediately and the module was retracted. After the go cue, in the interaction stage, the monkey released the start button, reached toward the module, and performed the corresponding hand interaction (see Behavioral Setup and Figure 1D for details). Successful completion marked the initiation of the reward stage and triggered the delivery of a small amount of water to the monkey. The monkey then returned its hand to the start button and pressed it again, triggering retraction of the module and defining the end of the trial.

Each session comprised 10 blocks, and each block contained one presentation of each of the 40 modules, yielding 400 trials per session and a total of 8,000 trials across the 20 recorded sessions (Figure 1B). A randomized module sequence was generated before the session and repeated in the same order across all 10 blocks. The monkey typically completed two approximately 1-h sessions per day, one in the morning and one in the afternoon.

### 4. Motion-capture system setup

To collect hand movement data without influencing the behavior of the subjects, we designed a markerless motion-capture system based on industrial vision cameras and machine learning keypoint recognition algorithms. Around the monkey’s workspace, eight cameras (acA1440-73gc, Basler) equipped with lenses (VS-LDA4, Vital Vision Technology) were arranged around the transparent operating table (Figure 2E). Logical identifiers ‘*Camera_0’* through ‘*Camera_7’* were assigned in ascending order of camera serial number. The cameras acquired BayerRG8 images at 1,440 × 1,080 pixels resolution. Exposure time was fixed at 2 ms and analog gain at 0. White-balance settings were adjusted separately for each camera to provide visually consistent color across views.

To keep the cameras synchronized, we used a microcontroller to generate trigger pulses. All cameras received the same rising-edge hardware trigger generated at a nominal frequency of 74 Hz with a 50% duty cycle. Images were demosaiced and stored as one Motion JPEG AVI file per camera. Acquisition used a modified version of JARVIS-AcquisitionTool 1.2 to support Basler cameras.^20^

Each recording contained eight AVI files, a frame-level metadata.csv file, and a triggerdata.csv file. The frame metadata retained logical camera identifiers, camera serial number, within-recording frame index, camera timestamp, and camera image identifier. The trigger file retained the trigger pulse index and associated digital-input states. These fields allow frame order, missing frames, and synchronization to be assessed independently of the nominal video frame rate.

### 5. Markerless hand-pose reconstruction

#### 5.1. Camera calibration

The eight-camera system was calibrated from synchronized videos of a standard checkerboard with 15-mm squares and an 11 × 8 array of internal corners. During recording, the checkerboard was moved throughout the recording space while its corners remained clearly visible in several camera views at the same time. Intrinsic parameters were estimated separately for each camera using Zhang’s calibration method,^21^ including focal length, principal point, and radial and tangential distortion coefficients. For extrinsic calibration, checkerboard corners visible simultaneously in multiple views were used in perspective-n-point and stereo-calibration procedures to estimate the rotation and translation of each camera in a common world coordinate system. Intrinsic and extrinsic calibration and subsequent optimization were performed with the calibration module of JARVIS-AnnotationTool.^22^ The resulting camera-specific parameters were saved as OpenCV YAML files. Each file contained the intrinsic matrix, lens-distortion coefficients, and relative camera pose used for image undistortion and multiview reconstruction. During the experiments, the intrinsic and extrinsic camera parameters were recalibrated from checkerboard recordings at intervals of approximately 3-4 days. To minimize calibration errors arising from the potential changes in camera position over time, inference for each experimental session used the calibration parameters estimated from the most recent calibration procedure.

#### 5.2. Keypoint annotation and quality control

Training and evaluation datasets were manually annotated in JARVIS-AnnotationTool^22^ using a hand-and-elbow skeleton with 24 keypoints comprising the elbow, the wrist center along with its medial and lateral points (labeled as Wrist_medial and Wrist_lateral in the dataset; defined relative to the pronated hand in the workspace, corresponding to the radial and ulnar styloid processes, respectively), and the carpometacarpal joint (CMC), metacarpophalangeal joint (MCP), interphalangeal joint (IP), as well as fingertip of each digit (Figure 2F). Subject-specific segment lengths were derived from CT data and used to constrain 3D reconstruction. The constraints covered the elbow-to-wrist segment and the wrist-to-fingertip chains through the CMC, MCP, and IP keypoints. Multiview annotations were accepted only if all reconstructed segment lengths were within 5 mm of the CT-derived values and the mean reprojection error of every keypoint across views was below 5 pixels.

Because the anatomical elbow was usually occluded by the primate chair, annotators marked a visible point along the continuation of the wrist-to-elbow direction in each view instead of attempting to localize the hidden joint directly. Multiview consistency and reprojection constraints were used to recover the corresponding 3D direction. The anatomical elbow position was then calculated from this direction and the CT-derived wrist-to-elbow length. The calculated elbow coordinate was used for model training and evaluation.

#### 5.3. Model training

A pose estimation model for monkey hand was trained with the JARVIS-HybridNet framework, which combines 2D and 3D convolutional neural networks for multiview 3D pose estimation.^23^ Human annotations served as ground truth for model training. The training set contained 1,179 synchronized eight-view frames (9,432 annotated images) across four sessions. These comprised two training sessions and two experimental sessions and covered all task types.

#### 5.4. Model evaluation

The evaluation set was drawn from an experimental session also represented in the training set. However, no frame appeared in both datasets. To cover the full behavioral paradigm, 40 blocks were identified using behavioral event markers, and two successful trials were selected from each block. For each trial, five synchronized frames were sampled at equal intervals between the start-button press and reward delivery. This procedure yielded 400 independently annotated eight-view frames.

To evaluate the quality of reconstruction, we calculated the error of 3D position for all the keypoints. For keypoint *i* in frame *j*, the Euclidean distance between the predicted and manually annotated 3D coordinates was calculated as:

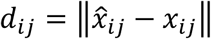

where *x̂_ij_* and *x_ij_* denote the predicted and manually annotated coordinates, respectively. Both sets of coordinates were reconstructed using the same intrinsic and extrinsic camera parameters. Therefore, no rigid rotation or translation was applied before error calculation. The 3D position error for keypoint *i* was then defined as the median of these distances across the *N* evaluation frames:

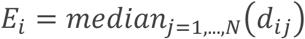

Quantitative model-performance results are reported in Technical Validation (Accuracy of markerless 3D hand-pose reconstruction model).

#### 5.5. Model inference and motion-capture preprocessing

The final monkey hand-pose estimation model trained and evaluated as described above (see Model training and Model evaluation) was deployed on a computing platform equipped with eight NVIDIA A100 GPUs with 80 GB of memory each (640 GB total GPU memory). Frame-by-frame multiview inference was performed jointly across the eight synchronized video streams acquired in each experimental session.

For each synchronized eight-view frame set, the model predicted the predefined hand and elbow keypoints and returned their 3D coordinates together with corresponding confidence scores. Because the model operates on individual frames independently, no temporal smoothing was applied during inference, and the predictions were kept as the direct frame-wise estimates.

We applied a multistep preprocessing pipeline to the model outputs to improve reconstruction quality and continuity. The preprocessing pipeline first identified abnormal frames. A frame was marked as invalid if any keypoint had a low confidence score (JARVIS output confidence^23^ < 0.1) or if an estimated bone length deviated by more than 35% from its CT-derived anatomical measurement. Missing and invalid data were reconstructed using Akima spline interpolation for gaps shorter than 200 ms, whereas longer gaps were left uninterpolated. A second bone-length check using the same threshold (> 35%) was then performed to discard frames containing excessive deviations introduced by interpolation.

For subsequent analyses, the processed coordinates were mapped into three reference frames: a world reference frame, with its origin and orientation defined by the ArUco marker on the primate chair; a wrist-global frame, with its origin translated to the wrist keypoint while retaining the global orientation; and a wrist-local frame, which retained the wrist origin but rotated the axes to align with the vector between the ulnar and radial wrist keypoints and the vector from the wrist keypoint to the base of the middle finger (MCP3). The wrist-local transformation removed the variation in global hand position and orientation.

Kinematic features were then extracted in the relevant reference frames and grouped into four categories: (1) joint angles, including MCP flexion-extension and abduction-adduction for digits 2-5, IP flexion-extension, and thumb carpometacarpal (CMC1) flexion and opposition; (2) pinch geometry, comprising the distances between the thumb tip and each of the other four fingertips; (3) global hand pose, including 3D Cartesian wrist position and hand orientation (pitch, roll, and yaw); and (4) dynamic features, including the linear velocities and accelerations of the wrist and fingertips. Finally, principal component analysis (PCA) was applied to the keypoint coordinates, and the first 20 principal components (PCs) were extracted for further analysis.

### 6. Electrophysiological recording and preprocessing

Electrophysiological signals were acquired with two parallel recording systems. Each system acquired data from a 1,024-channel electrode group through the SpikeGadgets Modular Stacking Headstage (MSH) datalogger system and Main Control Unit (MCU). For each system, a workstation ran Trodes 2.8.1 to save the recorded data from the MCU in the native .rec format. A complete session therefore contained 2,048 channels. Signals were sampled at 30 kHz and stored as signed 16-bit integers (int16), using a conversion gain of 0.195 μV per bit and an offset of 0 μV. MCU ticks, 64-bit system-clock values, and enabled digital inputs were retained as additional timing and event fields.

Each 1,024-channel electrode group contained eight physical 128-channel arrays. All preprocessing was performed separately for each array. The 128 channels corresponding to the dislodged array were retained in the released dataset but omitted from subsequent analyses except for the neural decoding benchmark. The original integer sample indices were retained throughout processing, and no temporal interpolation was applied. Threshold-crossing events (TX) and spike-band power (SBP) were computed for each array in a separate processing branch. The stored int16 values were converted to float32 voltages using the channel gains in the .rec header (0.195 μV per bit; zero offset). The signals were band-pass filtered at 250-5,000 Hz with a fourth-order Butterworth filter.^24^ The filter was implemented in second-order-section form and applied in both forward and reverse directions, followed by CMR within each array. Each recording comprised a single .rec file and was represented as a single SpikeInterface segment; therefore, segment_index was 0 throughout. For TX extraction, the threshold for each channel was set to −3.5 times the population standard deviation estimated from the first 10 s of the processed signal. A TX event was marked when a sample fell below this threshold after the preceding sample had been at or above it. Each crossing was stored with its zero-based sample index, zero-based channel index, segment_index (0), and amplitude at the crossing sample in μV. For SBP, one power value per channel was calculated for each successive 20-ms bin (600 samples at 30 kHz). Each value was the mean squared voltage within the bin and was expressed in μV². The bins did not overlap, and any incomplete final bin was discarded. Each bin was stored as one row containing the 128-channel power values, segment_index (0), the inclusive start sample index, the integer center sample index (start + 300), and the exclusive end sample index.

Processed data were stored in two complementary types of structured files. For each 1,024-channel electrode group, one synchronization file preserved timing and digital events, whereas eight array files stored array-specific information and results. A complete session therefore contained two synchronization files and 16 array files. In the synchronization file, each valid .rec packet, corresponding to one sample across channels, was represented by a clock-table row containing its 0-based integer sample index, raw MCU tick, and Unix system time in nanoseconds. The mapping between integer sample indices and Unix system time aligned electrophysiological samples to a common timeline used by the motion capture system and the behavioral program. The rising edges of TTL signals sent by the behavioral program were stored using the same sample-index scale. Each array file corresponded to one physical 128-channel array. Its channel map linked acquisition channels to electrode contacts and specified the relative coordinates of those contacts within the array. The array file also contained sorted units with quality annotations, TX events, and SBP values. These results used the same sample indices as the associated synchronization file, enabling direct temporal alignment. Source-file identifiers, software versions, and processing parameters were retained as provenance.

### 7. Cross-system temporal alignment

The cameras, behavioral workstation, and two electrophysiology workstations were synchronized through a common Precision Time Protocol (PTP) hierarchy. A Huawei S5735 switch served as the local grandmaster for the camera hardware clocks, and a second synchronized switch distributed the same time base to the workstations for behavioral program and electrophysiological recording. The system time of the workstation for video acquisition was not used for alignment. Native video-frame, electrophysiology-sample, spike, and behavioral-event indices were instead mapped to the PTP-derived timeline recorded by their respective modality. Because all modality-specific timelines were derived from the same grandmaster, all these modalities could be compared on a common timeline without altering the archived source data.

All experimental videos were recorded with camera PTP enabled. Camera timestamps therefore used a scale of 1 ns per tick and were stored as ‘*frame_timestamp’* in ‘*metadata.csv’*. Electrophysiology samples retained a mapping between their native 30-kHz sample index and the PTP-disciplined workstation system time. This mapping was exported by the *’exporttime’* utility as *’continuoustime.dat’* in the Trodes software. Behavioral event times were read from the ‘*timestamp’* field written using the PTP-disciplined workstation system time.

Camera synchronization was evaluated from corresponding frame timestamps across the eight views and from the intervals between consecutive frames. In the representative recording session, each camera contained 350,103 frames. The median cross-camera timestamp spread was 296 ns, the 95th percentile was 344 ns, and the maximum was 424 ns. The mean frame interval was approximately 13.509688 ms, corresponding to a measured acquisition rate of approximately 74.02095 Hz.

### 8. Neural decoding benchmark

The neural decoding benchmark was designed to test whether the released electrophysiological features retain information about three-dimensional hand kinematics. To separate the effects of model class from differences in data handling, all decoders used the same session-wise inputs, prediction targets, trial definitions, and train-test partitions. The benchmark included conventional linear and nonlinear baselines, state-space and recurrent models, a latent-dynamics model, and the proposed Latent Manifold-guided Lag-aware Fusion (LMLF) decoder (Figure 3B).

**Figure 3.**
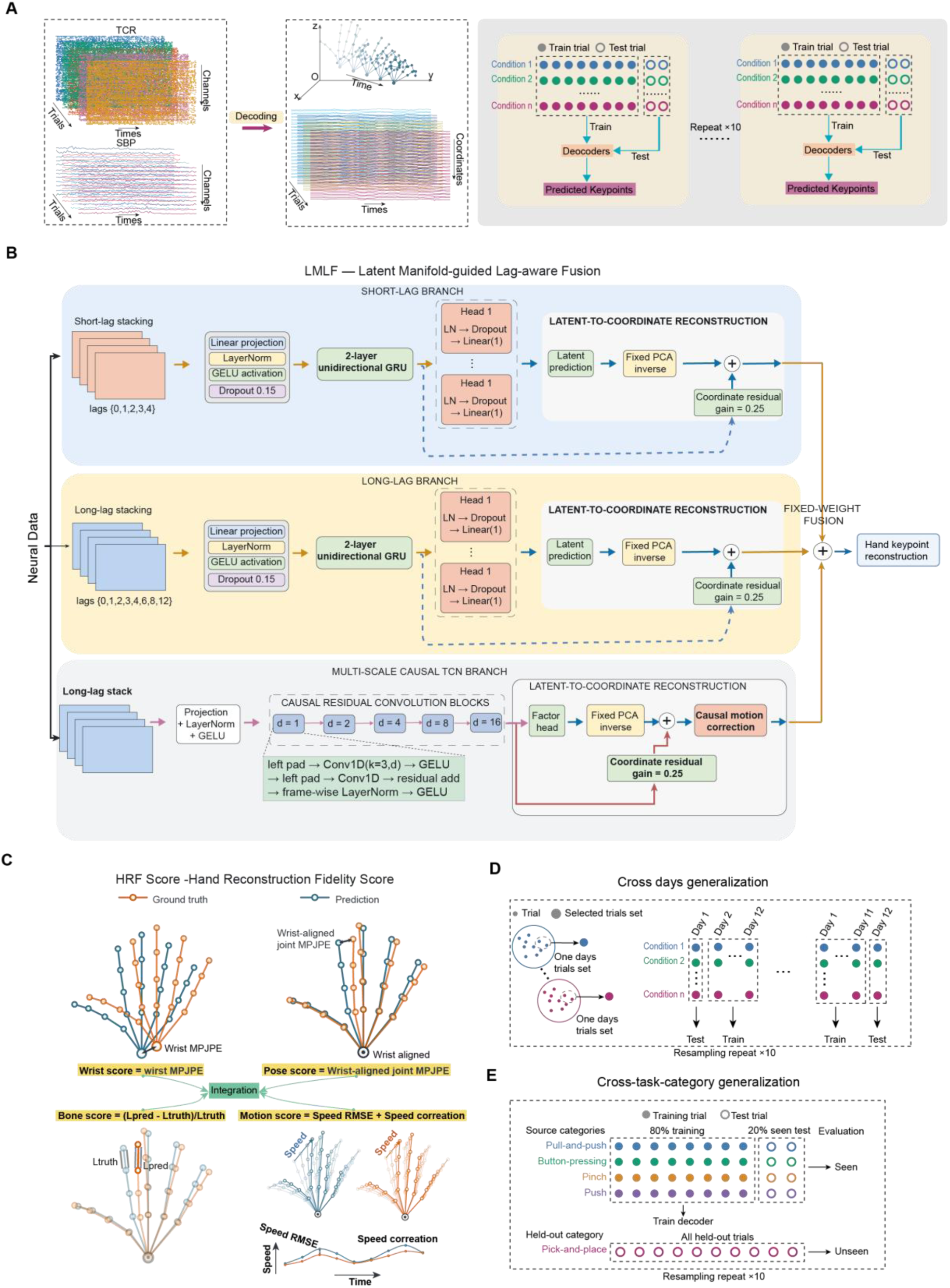
Hand movement decoding benchmark design and LMLF decoding strategy. (A) Neural inputs, predicted 3D hand keypoints, and the repeated condition-stratified 80/20 trial-splitting protocol. (B) LMLF architecture, comprising short-lag GRU, long-lag GRU, and multiscale causal temporal convolutional network branches followed by fixed-weight fusion. (C) Composition of the Hand Reconstruction Fidelity Score that covers global translation, wrist-centered pose, local motion, and skeletal consistency components. (D) Cross-day generalization test examined the generalization ability of algorithms over different experimental dates. All sessions from one of 12 recording days were held out for testing, using every valid trial from that day. Within each remaining source session, 20% of trials from each task condition were sampled for training, rounded up to retain at least one trial per condition. Source-trial sampling was repeated 10 times for each held-out day, while the test set remained fixed. Small dots represent individual trials, and large filled circles represent selected trial sets. (E) Task-category generalization test examined the generalization ability of algorithms over different task categories. One of the five task categories was held out as an unseen test category, while the remaining four categories formed the training folds. Within the training folds, 80% of trials were used for model fitting, and the remaining 20% were reserved as the seen test set. This 80/20 split was repeated 10 times for each held-out task category. Small dots represent individual trials, and large filled circles represent selected trial sets.

#### 8.1. Decoder inputs and prediction targets

At each video frame, the two 1,024-channel electrode groups contributed 2,048 TX features and 2,048 SBP features, which were concatenated into a 4,096-dimensional neural observation. The kinematic target comprised the wrist and four joints from each digit, yielding 21 keypoints and 63 Cartesian coordinates (**Figure 3**A). Elbow, Wrist_medial, and Wrist_lateral were retained in the released kinematic records but were not included in the decoding target. The corresponding file organization is described in Data Records.

Models were fitted separately for each recording session. Normalization, channel selection, dimensionality reduction, and parameter estimation were performed exclusively on the training trials of each split. The held-out trials were used only for inference and scoring. This separation prevented information from the test trajectories from influencing either the neural representation or the kinematic latent space.

#### 8.2. Trial construction and quality control

To enable a reliable and consistent evaluation of decoding algorithms, we segmented the continuous recordings into individual trials and retained only trials labelled as successful. For each retained trial, a common temporal window was extracted from the TX, SBP, and hand keypoint streams, spanning from 40 frames before trial onset to 40 frames after the reward delivery event. Trial onset marks the beginning of task execution, whereas the reward delivery event indicates when the subject completed the task. Together, these two events define the core task period. The additional frames before trial onset and after the reward delivery event provide temporal context surrounding task execution, allowing each segment to capture the complete hand trajectory associated with successful task performance as well as its immediately preceding and subsequent context.

Data quality control was applied separately to the core task period and the surrounding contextual periods. A trial was excluded if any NaN values were present in the keypoint coordinates, TX, or SBP data between trial onset and the reward delivery event, ensuring that every retained trial contained complete neural and kinematic measurements throughout task execution. In contrast, when NaN values occurred only within the contextual periods before trial onset or after the reward delivery event, the trial was retained and the corresponding boundary of the extraction window was shifted inward to exclude the invalid samples. As a result, the segmented trials may vary in duration, while each retained segment preserves a complete, finite-valued core task trajectory and maintains temporal alignment across the neural and kinematic data streams.

#### 8.3. Decoder comparison set

The comparison set spans linear, nonlinear, state-space, recurrent, and latent-dynamics approaches so that performance differences can be interpreted with respect to model capacity and temporal assumptions rather than a single algorithm family (Table 2). Linear and supervised latent baselines include Ridge regression, Elastic Net, partial least squares (PLS), and PCA-Ridge regression. Wiener Cascade adds a coordinate-wise polynomial nonlinearity to a causal linear filter, whereas support vector regression (SVR) provides a kernel baseline. Kalman filter represents linear dynamical state estimation. Simple recurrent neural network (RNN), gated recurrent unit (GRU), and long short-term memory (LSTM) decoders provide matched recurrent baselines for direct coordinate prediction. Finally, selected-channel latent factor analysis via dynamical systems (LFADS) with a linear readout evaluates a learned latent neural-dynamics representation.^25^ This coverage follows the broad distinction between static regression, dynamical filtering, and nonlinear sequence models used in neural decoding.^26^

**Table 2.** Decoder families included in the neural decoding benchmark.

| Family | Project methods | Benchmark role | Temporal treatment |
| --- | --- | --- | --- |
| Regularized linear | Ridge; Elastic Net | Low-variance and sparse linear references | Causal lagged features |
| Latent linear | PLS; PCA-Ridge | Linear low-dimensional representations | Causal lagged features |
| Nonlinear regression | Wiener Cascade; SVR | Polynomial and kernel nonlinear references | Causal history windows |
| State-space | Kalman Filter | Linear dynamical filtering reference | Recursive state update |
| Recurrent neural | Simple RNN; GRU; LSTM | Direct nonlinear sequence decoding | Unidirectional recurrence |
| Latent dynamics | Selected LFADS + Linear | Learned neural-dynamics representation | Sequential latent model |
| Proposed | LMLF | Latent kinematics with multiscale temporal fusion | Strictly causal multi-branch history |

#### 8.4. LMLF kinematic manifold and lag-aware channel selection

LMLF decomposes decoding into two coupled problems (Table 3). Estimating a compact kinematic state from neural history and reconstructing the complete hand pose from that state. This design avoids treating the 63 output coordinates as unrelated targets. Low-dimensional population states have previously been used to preserve movement-related information in motor-cortical decoding,^27^ and full-hand kinematics can be represented by compact latent variables.^28^ In LMLF, principal component analysis (PCA) supplies a deterministic and reproducible kinematic manifold. PCA is used as a linear structural constraint rather than as a claim that hand kinematics are intrinsically linear. Nonlinear autoencoders can yield lower kinematic reconstruction errors in other hand-movement datasets.^28^

**Table 3.** LMLF architecture and optimization settings.

| Component | Setting | Function |
| --- | --- | --- |
| Input | 2,048 TX + 2,048 SBP features per frame | Joint neural observation |
| Preprocessing | Training-partition normalization; Gaussian $\sigma = 2$ frames | Scale alignment and temporal denoising |
| Kinematic manifold | 32 PCA factors | Compact shared output representation |
| Channel selection | 320 channels; top three factor correlations; lags 0-12 | Lag-sensitive neural feature selection |
| Short GRU | 2 layers; hidden 256; lags 0-4 | Immediate and short-range dependencies |
| Long GRU | 2 layers; hidden 256; lags 0,1,2,3,4,6,8,12 | Delayed recurrent dependencies |
| TCN | Hidden 320; kernel 3; dilations 1,2,4,8,16 | Finite multiscale causal context |
| Branch output | 32 factors + inverse PCA + residual gain 0.25 | Manifold reconstruction with coordinate correction |
| Objective weights | Coordinate 1.0; factor 0.2; velocity 0.1 | Joint pose, latent, and motion supervision |
| Optimization | 120 epochs; batch 8; AdamW; learning rate $10^{-3}$; weight decay $10^{-4}$ | Fixed training configuration |
| Fusion | 0.36 short GRU; 0.24 long GRU; 0.40 TCN | Combine complementary temporal estimates |

The 63-dimensional training-partition keypoint coordinates are standardized and decomposed by PCA. The first 32 components define the kinematic factors used as branch-level supervision. Both the standardization parameters and PCA basis are stored with the session checkpoint and reused unchanged during test-partition inference. This separation prevents test trajectories from influencing the latent coordinate system.

Channel selection is explicitly conditioned on temporal lag. For each neural channel, absolute correlations with the 32 training-partition PCA factors are evaluated at lags {0, 1, 2, 3, 4, 6, 8, 12}. At each lag, the three largest factor correlations are averaged, and the maximum of these lag-specific scores becomes the channel score. The 320 highest-scoring channels are retained. This criterion favors channels that are informative for at least a subset of the kinematic manifold while allowing their strongest association to occur at different causal delays. Only current and preceding neural frames are used.

#### 8.5. LMLF multiscale temporal encoding and coordinate readout

After training-partition normalization and Gaussian smoothing with a standard deviation of two frames, a lag-aware selection stage retained 320 neural channels. The selected features were supplied to three parallel causal encoders: a short-context GRU, a long-context GRU, and a dilated temporal convolutional network (TCN). Each encoder predicted the same 32-dimensional PCA representation together with a small coordinate-space residual. The three reconstructed poses were subsequently combined using fixed fusion weights, so that the kinematic manifold constrained the output structure while the temporal branches captured complementary short- and long-range dependencies.

The short-context GRU branch receives explicit lags {0, 1, 2, 3, 4}, and the long-context GRU branch receives {0, 1, 2, 3, 4, 6, 8, 12}. Both inputs are projected to 320 dimensions and processed by two-layer unidirectional GRUs with 256 hidden units. GRU gating provides adaptive recurrent memory with fewer gating operations than LSTM.^29^ The two GRU branches are not parameter-shared, permitting the short branch to specialize in local changes and the long branch to represent delayed dependencies.

The TCN branch uses the long-lag input, a 320-dimensional projection, and five causal residual blocks with kernel size 3 and dilations {1, 2, 4, 8, 16}. Each block contains two left-padded convolutions. Consequently, the convolutional stack has a 125-frame receptive field, and the explicit 12-frame lag extends the maximum available past support to 137 frames. Dilated causal convolutions provide a finite multiscale history that is complementary to recurrent state propagation.^30^ Layer normalization is applied independently within each frame. Normalization across time is deliberately avoided because it would allow future samples to alter earlier representations.

Each GRU branch uses 32 separate scalar readout heads. The heads share the branch temporal representation but do not share their terminal linear parameters, allowing different PCA factors to use distinct mappings from the same neural state. The TCN branch uses a 32-dimensional factor head. In every branch, predicted standardized factors are transformed to keypoint coordinates using the fixed inverse PCA mapping. A zero-initialized 63-dimensional residual head, multiplied by 0.25, corrects components that are not represented adequately by the linear manifold. Zero initialization makes the initial branch prediction coincide with manifold reconstruction and introduces coordinate corrections gradually during optimization.

#### 8.6. LMLF optimization, fusion, and inference

Each temporal branch is optimized independently with a coordinate loss, a latent-factor loss, and a first-difference velocity loss. Let Y and Y-hat denote the ground-truth and reconstructed coordinates, Z and Z-hat the corresponding PCA factors, and Delta the first-order temporal difference operator. The branch loss is:

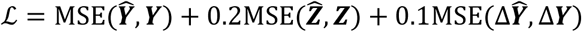

The coordinate term is dominant; the factor and velocity terms have weights 0.2 and 0.1, respectively. Each branch is trained for 120 epochs with AdamW, a learning rate of 10⁻³, weight decay of 10⁻⁴, mini-batches of eight trials, dropout of 0.15, and gradient-norm clipping at 1.0. Training uses seed 42.

During inference, the three coordinate predictions are combined using fixed weights. If Y-hat-short, Y-hat-long, and Y-hat-TCN denote the short-GRU, long-GRU, and TCN outputs, respectively, the fused prediction is:

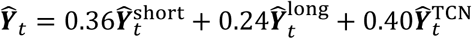

A causal motion correction is applied to the TCN prediction before fusion. It uses only the current prediction and the immediately preceding output, with beta = -0.5 and motion scale = 4.0. It therefore introduces no future context. The fixed fusion weights place 0.36 on the short GRU, 0.24 on the long GRU, and 0.40 on the TCN. In the present implementation, the three branches are trained independently, and the fusion weights are specified in the configuration rather than estimated from test data.

### 9. Neural decoding evaluation

Neural decoding was evaluated at two complementary levels. Held-out reconstruction metrics quantified the kinematic information retained in the electrophysiological features, whereas implementation-level tests assessed checkpoint reproducibility and strict temporal causality. The same evaluation protocol was applied to every decoder family to avoid method-specific differences in data selection or aggregation.

#### 9.1. Train-test partitioning and held-out evaluation

To maintain a balanced representation of experimental conditions across the training and test sets, we adopted a repeated condition-stratified random splitting strategy. Trials were first grouped according to their experimental conditions for each session. Within each condition, approximately 20% of the trials were randomly assigned to the test set, while the remaining 80% were assigned to the training set. Trials selected from all conditions were then pooled to construct the final test set, with the remaining trials forming the corresponding training set. This procedure was independently repeated 10 times to generate 10 train-test splits (**Figure 3**A), while preserving the representation of different experimental conditions in each split whenever possible. For conditions containing too few trials to support an 80/20 split, priority was given to assigning trials to the test set to ensure their representation in the evaluation data. The same set of 10 data splits was subsequently used for all decoding methods to ensure consistent comparisons across models.

For every partition, each algorithm was fitted independently within each session and evaluated only on the held-out trials. Preprocessing statistics and model parameters were not refitted during evaluation. Trial-level quantities were first aggregated within sessions, and benchmark-level values were then calculated as equally weighted means across the predefined set of 20 sessions. This two-stage aggregation prevented sessions with more or longer trials from dominating the reported values.

Reconstruction quality was summarized using the Hand Reconstruction Fidelity Score (HRFS), which separates global translation, wrist-centered pose, local motion, and skeletal consistency. Five conventional metrics were reported alongside HRFS to retain direct comparisons with standard neural-decoding analyses.

#### 9.2. Hand Reconstruction Fidelity Score

The Hand Reconstruction Fidelity Score (HRFS) is a composite measure of agreement between reconstructed and ground-truth hand movements. It evaluates four complementary properties of each predicted trajectory: global translation, wrist-centered pose, local motion, and skeletal consistency. This decomposition separates translation of the hand from finger articulation while retaining explicit measures of temporal dynamics and bone-length preservation (**Figure 3**C).

For a trial containing T frames and J = 21 keypoints, let the unaccented and hatted vectors in the equations below denote the ground-truth and predicted three-dimensional coordinates of keypoint j at frame t, respectively, and let w denote the wrist. Each component is evaluated only where the required ground-truth samples are finite. Its coverage cₖ is the fraction of these ground-truth-valid samples for which the corresponding prediction is also finite. Coverage therefore enters the component score directly rather than allowing missing predictions to be ignored. For compactness, [*z*]^100^ denotes min(100, max(0, z)) in the equations. For all evaluation metrics, we include only samples for which both the ground-truth and predicted coordinates are valid. Wrist reconstruction fidelity is quantified as the mean Euclidean distance between the predicted and ground-truth wrist positions, reflecting the accuracy with which the model reconstructs the global spatial position of the hand. To evaluate hand pose independent of the global translation of the hand, each non-wrist keypoint is expressed relative to the wrist position in the same frame. We then compute the mean Euclidean distance between the predicted and ground-truth wrist-centered coordinates, thereby measuring the accuracy of the reconstructed finger and joint configuration. The corresponding static component scores are:

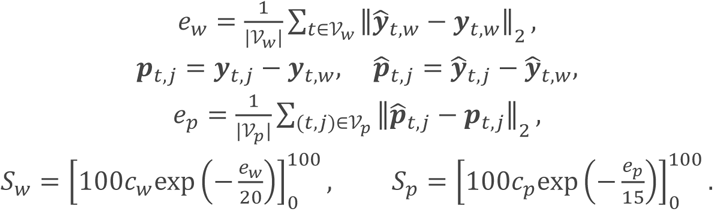

Motion fidelity is computed from first-order finite differences of the wrist-centered trajectories. The sampling interval Δt is the median positive finite difference between consecutive frame timestamps, with 1/30 s used only when no valid timestamp difference is available. The velocity error is the root mean squared error over all coordinates of non-wrist keypoint pairs in valid adjacent frames. The velocity-correlation term is the mean Pearson correlation computed separately for each non-wrist keypoint-coordinate velocity series. Series with fewer than two samples are omitted. Identical constant series contribute a correlation of one, whereas a series is omitted when only one member of the pair is constant. If no eligible series remains, the correlation term is set to zero. The motion component is scored as:

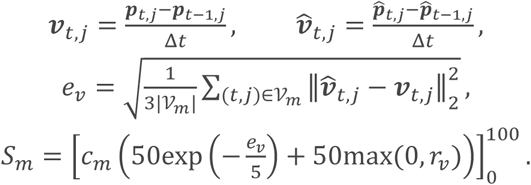

Skeletal consistency is evaluated on the 20 edges of the predefined hand skeleton, comprising the four consecutive segments of each of the five wrist-to-fingertip chains. For every valid frame-edge pair, the corresponding ground-truth and predicted bone lengths are computed from the parent and child coordinates; p(e) and c(e) denote the parent and child indices of edge e, respectively. The bone error is the mean absolute relative length error; a denominator floor of 10⁻⁸ is used for numerical stability. The bone component is scored as:

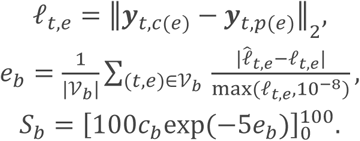

All four component scores are clipped to the interval [0, 100]. The exponential mappings use scale constants of 20 for wrist-position error, 15 for local-pose error, and 5 for velocity error; the bone penalty coefficient is 5. The first three constants are expressed in the corresponding position or velocity units of the input kinematics, whereas the bone error is dimensionless. For a component with a defined ground-truth denominator, zero coverage or a non-finite error produces a score of zero. A component is unavailable only when its coverage denominator cannot be defined.

The trial-level HRFS is the weighted mean of the available component scores. The base weights are 0.25 for Wrist, 0.35 for Pose, 0.20 for Motion, and 0.10 for Bone. The remaining 0.10 weight is reserved for cross-session stability (not evaluated here). When one or more component scores are unavailable, the remaining weights are renormalized by their sum. The set A in the equation contains the available components. The trial-level HRFS is:

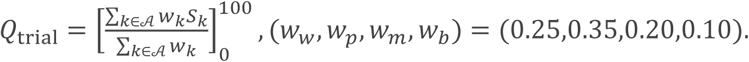

Within a session, trial scores are averaged with equal weight and clipped to [0, 100]. The benchmark-level HRFS is then the equally weighted mean of the session scores, also clipped to [0, 100], where N is the number of trials in a session and M is the number of evaluated sessions. The session and benchmark scores are:

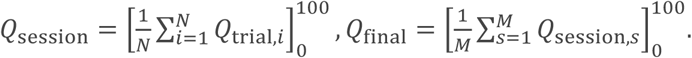

A complete, exact reconstruction with full coverage receives 100 for every component and at every aggregation level. Higher HRFS values therefore indicate closer agreement in hand location, wrist-centered configuration, local velocity, and bone length. The metric does not include a latency bonus or penalty; runtime limits are enforced separately by the evaluation protocol.

#### 9.3. Conventional reconstruction metrics

In addition to the proposed Hand Reconstruction Fidelity Score (HRFS), we report five conventional reconstruction metrics: the macro-averaged coefficient of determination (Macro R²), macro-averaged Pearson correlation coefficient (Macro Pearson r), root mean squared error (RMSE), mean absolute error (MAE), and mean per-joint position error (MPJPE). These metrics complement HRFS by separately reporting coordinate-wise goodness of fit, trajectory correlation, scalar coordinate error, and three-dimensional joint-position error.

For each algorithm and session, the ground-truth and predicted trajectories from all held-out trials are concatenated along the frame axis before evaluation, producing arrays with F frames, 21 keypoints, and three Cartesian coordinates per keypoint. The arrays are evaluated on the reconstructed coordinate scale; the metric implementation does not apply additional standardization. Every calculation uses a pairwise finite mask, so only entries for which both the ground truth and prediction are finite contribute to the corresponding metric.

For Macro R², each session array is reshaped into 63 coordinate dimensions, corresponding to the three coordinates of each of the 21 keypoints. A separate coefficient of determination is computed for coordinate dimension d over its jointly finite frames, and the session-level Macro R² is the unweighted mean across eligible dimensions. Macro R² is defined as:

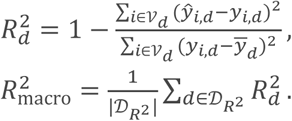

Here, the mean ground-truth value for dimension d is computed over its valid set. A dimension is included only when it contains at least two jointly finite samples and its ground-truth sum of squares exceeds 10⁻¹². The coordinate-level and macro-averaged R² values are not clipped. Negative values are retained.

Macro Pearson r is calculated using the same 63 coordinate dimensions and pairwise-valid frames. Pearson correlation is computed separately for each eligible dimension, after which the coefficients are averaged with equal weight. Macro Pearson r is defined as:

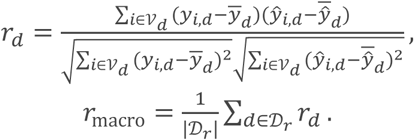

A coordinate dimension contributes to Macro Pearson r only when it contains at least two jointly finite samples and both its ground-truth and predicted standard deviations exceed 10⁻¹². Constant ground-truth or predicted dimensions are therefore excluded from the macro average rather than assigned an arbitrary correlation value.

RMSE and MAE quantify scalar coordinate error. Let Ωc contain all jointly finite frame-keypoint-coordinate entries within a session. Unlike the macro metrics, which first assign equal weight to eligible coordinate dimensions, RMSE and MAE pool all valid scalar coordinates before averaging. They are computed as:

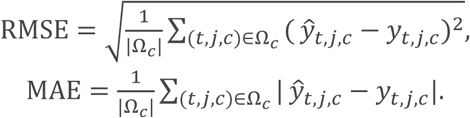

RMSE applies a quadratic penalty to the coordinate residuals, whereas MAE averages their absolute magnitudes. Both metrics retain the units of the keypoint coordinates, and lower values indicate smaller reconstruction errors.

MPJPE evaluates three-dimensional joint displacement. A frame-keypoint pair is valid only when all three ground-truth coordinates and all three predicted coordinates are finite. Let Ωj denote the resulting set of valid pairs. MPJPE is computed as:

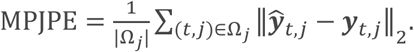

MPJPE averages the Euclidean distance between corresponding reconstructed and ground-truth keypoints. It has the same spatial unit as the keypoint coordinates, and lower values indicate closer joint-position agreement.

All five metrics are first computed independently for each session. The reported benchmark value for metric m is the mean of the corresponding session-level values over the predefined set of 20 sessions:

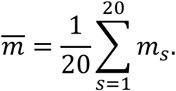

This aggregation gives every session equal influence on the reported result, irrespective of the number or duration of its held-out trials. For Macro R² and Macro Pearson r, higher values indicate better performance. For RMSE, MAE, and MPJPE, lower values indicate better performance.

#### 9.4. Checkpoint reproducibility and causal integrity

LMLF was trained independently within each session from the TX and SBP features and the aligned kinematic targets. Each checkpoint stores the neural-feature statistics, selected-channel indices and scores, target standardization parameters, PCA parameters, three branch state dictionaries, and the training configuration. Checkpoint round-trip tests verified that saving and reloading these components reproduced test-partition inference.

Strict causality was tested by perturbing neural samples after a selected time point and comparing the prediction prefix before and after perturbation. The prefix was required to remain unchanged within an absolute tolerance of 10⁻⁶. This test jointly covered explicit lag construction, recurrent directionality, convolutional padding, and per-frame normalization. It verifies model-level causality but does not measure streaming latency, buffer management, or hardware throughput. Because the present configuration uses one training seed per split, comparisons capture variability across sessions and data partitions but not between-seed optimization variability.

#### 9.5. Cross-day generalization

We evaluated cross-day decoding generalization performance of selected algorithms. To reduce the differences in keypoint representation caused by session-specific camera coordinate systems, we aligned the three-dimensional hand keypoints before training and evaluation. Session-specific calibration transformed the keypoints into the world reference frame defined by the ArUco marker on the primate chair, providing a consistent spatial reference across recording days. Each keypoint was translated relative to the marker origin and projected onto its orthonormal axes, with a consistent z-axis convention. Then we used a leave-one-recording-day-out protocol across 20 sessions from 12 recording days (**Figure 3**D). In each fold, all valid trials from one recording day were reserved for testing. Within each remaining session, 20% of trials from each task condition were sampled without replacement for training, rounded upwards to retain at least one trial per condition. Sampling was repeated ten times per held-out day, with identical training selections and fixed test sets across decoders. All preprocessing and model fitting used only the selected source-training data, without target-day adaptation.

Using these shared splits, we evaluated PCA-Ridge, LSTM, and LMLF with their previously described architectures unchanged. Unlike within-session decoding, each model was trained jointly on selected trials from multiple source sessions and evaluated on an unseen recording day. Normalization, feature selection, and PCA fitting, where applicable, used only the selected source-training data. For LMLF, feature-selection ranks were averaged across source sessions. Session-balanced trial sampling was used during training to limit unequal session contributions. LSTM and PCA-Ridge retained their respective fitting procedures on the pooled source data.

#### 9.6. Cross-task generalization

To evaluate the generalization capability of the neural decoding model across different task categories, we adopted a leave-one-task-category-out training and testing strategy based on the neural decoding benchmark described above. In each experiment, one of the five complete task categories was held out as the unseen category, while the remaining four task categories were used to construct the training set. Each of the five task categories was held out in turn to evaluate the decoding performance when transferring from seen task categories to an unseen task category.

As shown in **Figure 3**E, for each held-out task category, all trials from that category were first assigned to the unseen test set. These data were not used for model training or for estimating any training-related parameters, and were reserved exclusively for final performance evaluation. The remaining four task categories constituted the training domain and were split according to the specific task conditions using an 80/20 partition. Specifically, 80% of the training-domain data were used for model fitting, while the remaining 20% served as an in-domain reference test set, denoted as seen.

For each Session and each held-out task category, the 80/20 data split was repeated 10 times. Different random seeds were used for the data partitions, while the model-training random seed was fixed at 42. An independent model was retrained for each split and used to generate predictions on both the seen test set and the complete unseen task category. The experiment covered 20 Sessions, 5 held-out task categories, and 10 repetitions for each split, resulting in1000 model-training runs. This experiment used only LMLF to evaluate cross-task-category generalization. Apart from the training/testing data-partitioning strategy, no additional modifications were made to the model architecture or the primary training parameters.

For each experiment, all training-related operations were performed exclusively using the 80% training-domain data, including data normalization, feature selection, and the fitting of PCA or other target-transformation parameters. The fully held-out unseen task category was not involved in any of these procedures and was not used for model training, early stopping, or hyperparameter selection. The 20% seen data within the training domain were used solely to evaluate performance on seen task categories and were not involved in model fitting. This design makes it possible to distinguish the model’s decoding ability within known task categories from its generalization ability to a completely unseen task category.

Model performance was evaluated exclusively using the HRFS. For each session and held-out task category, the HRFSs on the seen and unseen test sets were calculated separately, and the results from the 10 repeated experiments were summarized. To quantitatively measure the differences in generalization performance among the task categories, we calculated the relative generalization gap, which is defined as:

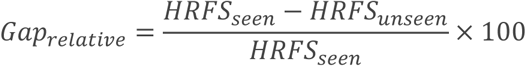

#### 9.7. Channel-availability sensitivity analysis

We performed a channel-availability sensitivity analysis to quantify how neural decoding changed as the number of available intracortical channels decreased. The dislodged eg1-8 array was excluded in every condition, leaving 1,920 eligible channels from the 2,048 implanted channels. Each retained channel contributed one TX feature and one SBP feature, so a condition with N physical channels contained 2N neural input features.

The unmasked condition contained all 1,920 eligible channels. The masked conditions retained 1,918, 1,916, 1,912, 1,904, 1,889, 1,858, 1,797, 1,676, 1,536, 1,434, 1,280, 1,120, 954, 800, 640, 512, 384, 256, 192, 128, 96, 80, 64, 48, or 32 physical channels. For each of 50 mask identifiers, a complete random permutation of the 1,920 eligible channels was generated using one fixed master seed together with that identifier. At a removal level of k channels, the channels indexed by the first k entries of the corresponding permutation were excluded. This procedure produced nested removal sets across channel counts. The fixed seed and mask identifiers allowed every mask to be regenerated exactly. Identical masks were applied across all 20 recording sessions and three decoders.

We evaluated LSTM, PCA-Ridge regression, and LMLF using their prespecified benchmark configurations. For LSTM, the remaining TX and SBP features were ranked using the root mean square of their correlations with the 63 target coordinates in the training data, and at most 160 features were retained. LSTM therefore used min(160, 2N) input features in a condition containing N physical channels. PCA-Ridge performed PCA on the masked training features and retained at most 64 principal components. LMLF used its lag-aware correlation criterion to retain at most 320 TX and SBP features and used all available features when fewer than 320 remained. Because TX and SBP from the same physical channel were ranked separately, the selected feature count was not necessarily the number of unique physical channels used by the neural models.

All three decoders used the same prediction targets and trial definitions. Every channel count, mask, session, and decoder used the first prespecified condition-stratified 80/20 train-test partition (repeat 0; split seed 42000) and training seed 42. We repeated feature normalization, feature selection, dimensionality reduction, and model fitting using only the masked training data, and evaluated performance on the corresponding held-out trials. The complete analysis comprised 60 unmasked fits and 75,000 masked fits (25 masked channel counts × 50 masks × 20 sessions × 3 decoders), for a total of 75,060 model fits.

We evaluated decoder performance using HRFS, Macro R², Macro Pearson r, RMSE, MAE, and MPJPE as defined above. For each decoder and channel count, the plotted value is the mean across sessions and masks. We estimated 95% confidence intervals using 10,000 two-way bootstrap samples that independently resampled the 20 sessions and 50 masks with replacement. The unmasked condition contained one configuration and was therefore resampled across sessions only.

## Data Records

The current release provides example trials of selected modules from the session-organized, decoding-ready dataset that aligns neural features, reconstructed hand kinematics, and trial metadata to a shared frame index. Raw videos, raw electrophysiological recordings, and other intermediate data products are not included in the present release. Every session directory follows the same six-file layout:

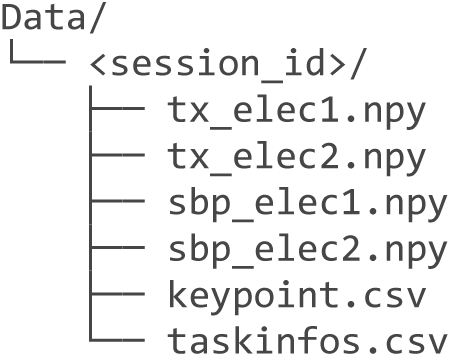

The four NumPy arrays store the TX and SBP feature families for the two electrode groups. Each array has shape T × 1,024 and data type float32, where T is the number of frames in the session. Concatenating the two electrode groups within a feature family produces a T × 2,048 matrix; concatenating TX and SBP yields the T × 4,096 neural representations used by the benchmark (Table 4).

**Table 4.** Decoding-ready files provided for each recording session.

| Field | Dimensions | Description |
| --- | --- | --- |
| tx_elec1.npy; tx_elec2.npy | $2 \times (T \times 1,024)$ | Frame-aligned TX features from electrode groups 1 and 2; float32. |
| sbp_elec1.npy; sbp_elec2.npy | $2 \times (T \times 1,024)$ | Frame-aligned SBP features from electrode groups 1 and 2; float32. |
| keypoint.csv | $T \times 73$ | Frame timestamp and x/y/z coordinates for 24 tracked keypoints. |
| taskinfos.csv | $N \times 40$ | Trial identifiers, labels, event times, frame indices, and alignment metadata. |

## Technical Validation

### 1. Active electrode channels during chronic implantation

The 16 electrode arrays were connected to the acquisition system as two electrode groups (abbreviated eg), denoted eg1 and eg2. Individual arrays were identified by their group and within-group position (e.g., eg1-1). Arrays eg1-1 through eg1-4 were implanted in PMv, whereas eg1-5 through eg1-8 targeted M1. Arrays eg2-1 through eg2-4 were implanted in AIP, and eg2-5 through eg2-8 were implanted in M1. The final M1 array, eg1-8, was dislodged from the cortical tissue during surgery. Array eg1-8 was therefore excluded, and a total of 1,920 channels from the remaining 15 arrays were used for analysis.

To assess the long-term performance of the implanted arrays, short electrophysiological recordings of approximately 15 min were obtained from Monkey Q while the animal was at rest. Recordings were performed approximately twice per week. After each recording, the signal from every channel was inspected visually by experienced technicians. A channel was classified as active when clear neural spiking activity was observed. We used data from 30 recording sessions spanning post-implantation days 1-120 and obtained 450 array-session observations.

The number of active channels varied substantially among arrays and across recording sessions (**Figure 4**A). Across the 15 implanted arrays, the mean was 8.53 ± 5.72 active channels per array on day 1, reached a transient maximum of 34.07 ± 21.04 on day 30, and was 7.07 ± 7.08 on day 120 (mean ± SD; **Figure 4**B). At the final time point, 10 of the 15 arrays retained at least one active channel, with 106 active channels in total. Across all 450 array-session observations, the mean was 13.21 ± 14.07 active channels per array, and the median was 9 channels. These measurements demonstrate that channels with detectable neural activity remained available in a subset of the implanted arrays for at least 120 days. However, the large dispersion between arrays and session-to-session variation indicate that active-channel availability was not uniform across the implant.

**Figure 4.**
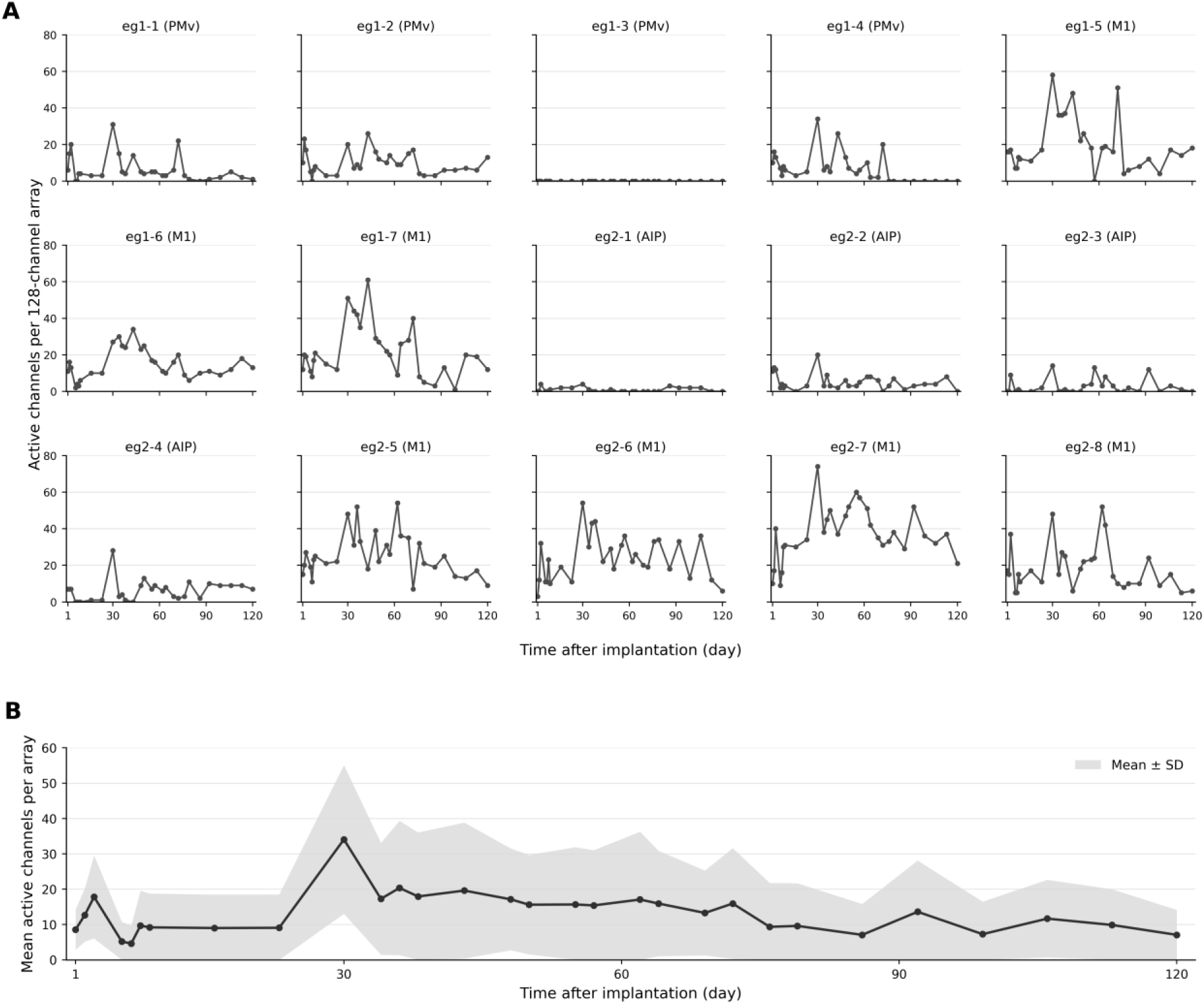
Availability of active electrode channels through the post-implantation period in Monkey. **Q.** (A) Active-channel counts for each of the 15 implanted 128-channel arrays. Panel titles give the array identifier and implantation area. Points denote the 30 recorded sessions from post-implantation days 1-120. Lines connecting observations are for visual guidance only. (B) Arithmetic mean active-channel count across the 15 implanted arrays. Shading denotes ±1 SD across arrays and is clipped at zero because channel counts cannot be negative.

### 2. Frequency-domain analysis of intracortical recordings

Frequency-domain analysis used the same resting-state recordings collected repeatedly from the implanted arrays for the active-channel assessment. The central 100 s of each recording was converted to μV and divided into epochs of 2 s. Constant or saturated channels were excluded, common-median referencing was performed within each 128-channel array, and artifact-contaminated epochs were rejected. An array-session observation was retained when at least 80 s of valid data remained.

Signals were resampled to 1 kHz, and channel-level power spectral density (PSD) was estimated using Welch’s method with 4.096-s Hamming windows and 50% overlap. Absolute power was calculated for delta (1-4 Hz), theta (4-8 Hz), alpha (8-13 Hz), beta (13-30 Hz), low gamma (30-45 Hz), and high gamma (70-150 Hz), excluding narrow regions around 50-Hz harmonics. We additionally calculated array-referenced 250-5,000 Hz root mean square (RMS) and the residual 50-Hz peak relative to its local spectral background.

Metrics were summarized within each physical array. Change during chronic implantation was estimated using a Theil-Sen slope for each of the 15 implanted arrays and reported as the median slope in dB per 30 days with a 95% confidence interval (CI) calculated through array-level bootstrap. The analyzed recordings retained a broad intracortical spectral profile with superimposed peaks at 50 Hz and its harmonics (**Figure 5**A). The headstage grounding configuration was changed to a single-point ground on post-implantation day 100 and was likely a major contributor to the reduction in high-gamma power in the late-period spectrum, although part of the offset was already evident before the change. The frequency-specific slopes nevertheless did not support describing all components as invariant. The median array-level slope was 0.52 dB/30 days for delta (95% bootstrap CI, 0.08 to 0.93), 0.35 dB/30 days for theta (−0.06 to 0.66), and 0.01 dB/30 days for alpha (−0.10 to 0.62). Higher-frequency power showed modest negative slopes: −0.33 dB/30 days for beta (−0.50 to −0.20), −0.28 dB/30 days for low gamma (−0.87 to −0.16), and −0.29 dB/30 days for high gamma (−0.79 to −0.08; **Figure 5**B). Thus, changes during chronic implantation were frequency specific rather than a uniform broadband loss.

**Figure 5.**
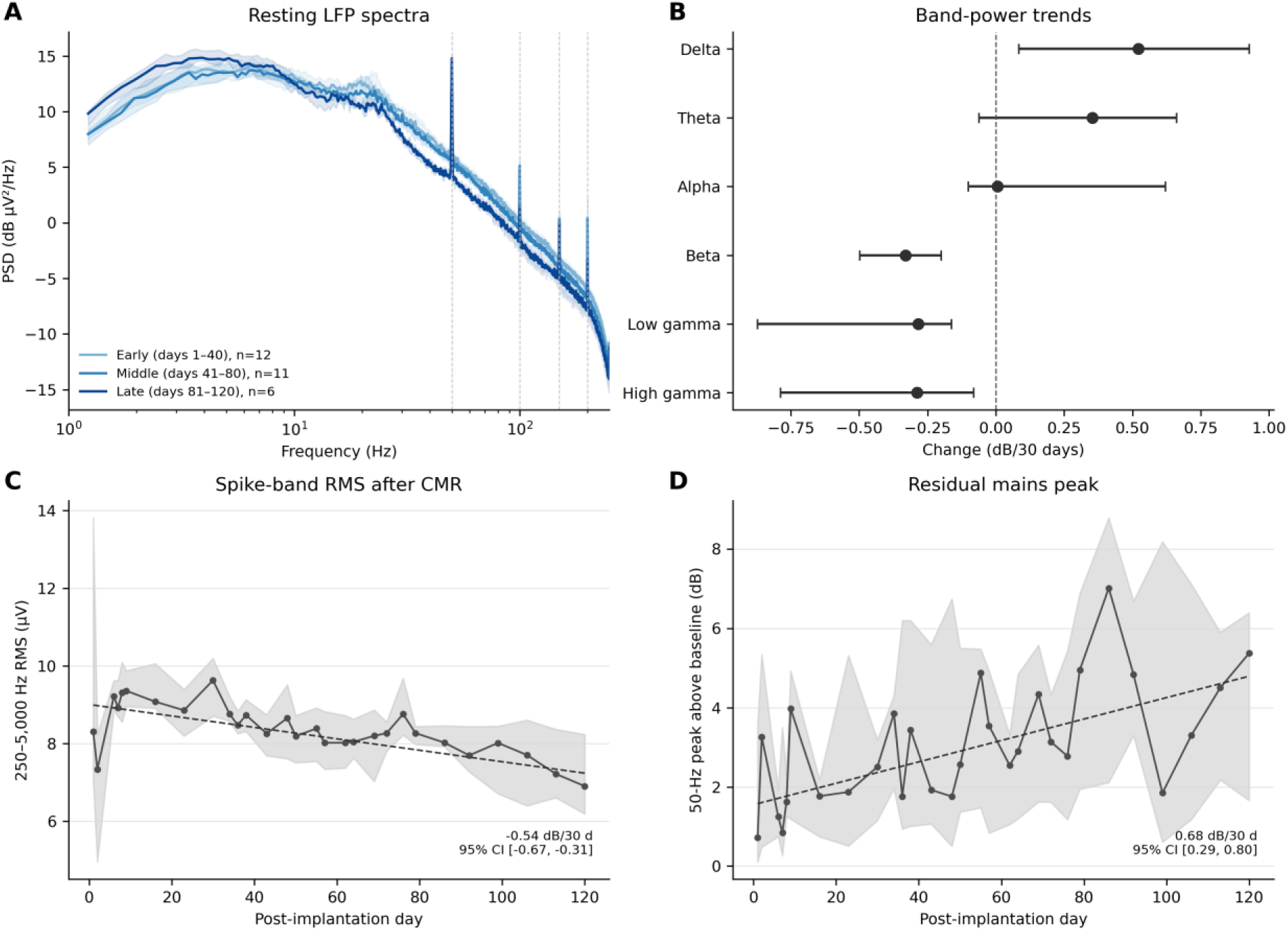
Resting-state spectra and recording-quality metrics in Monkey. **Q.** (A) Resting-state PSD during the early (days 1-40; n=12 sessions), middle (days 41-80; n=11), and late (days 81-120; n=6) periods. Within each session, PSD curves were summarized across the 15 implanted arrays. Lines denote the median across sessions in each period, shading denotes the interquartile range across sessions, and vertical dashed lines mark 50-Hz harmonics. (B) Median array-specific Theil-Sen slope and array-level bootstrap 95% confidence interval for each frequency band. (C) Session median and interquartile range of 250-5,000 Hz RMS after array-wise CMR. (D) Residual 50-Hz peak relative to its local spectral baseline. Dashed trend lines in C and D are robust Theil-Sen fits.

Spike-band RMS after CMR decreased from a cross-array median of 8.31 μV on day 1 to 6.90 μV on day 120, with a median slope of −0.54 dB/30 days (95% bootstrap CI, −0.67 to −0.31; **Figure 5**C), providing no evidence of a progressive broadband noise-floor increase. In contrast, the residual 50-Hz peak increased from 0.72 dB above its local baseline on day 1 to 5.38 dB on day 120. Its median slope during chronic implantation was 0.68 dB/30 days (0.29 to 0.80; **Figure 5**D). The data therefore retained measurable broadband and band-limited signals through day 120, but the increased mains peak and modest beta/gamma reductions should be considered in downstream spectral analyses. These findings complement, rather than replace, the manual active-channel assessment because resting band power is influenced by neural state as well as by the electrode-tissue interface.

### 3. Task-related modulation of intracortical field potentials

Functional signal content was evaluated in twelve representative standardized task sessions on post-implantation days 15, 29, 36, 48, 55, 62, 70, 79, 86, 99, 107, and 115. These sessions were distributed across the chronic implantation period and each contained 400 trials. Sixty successful trials per session were selected with balanced sampling across task identities. Behavioral go cue times were mapped to electrophysiological sample indices using PTP-derived *’continuoustime.dat’* files. This event was used as an explicit task-alignment marker and was not labelled as kinematically determined movement onset.

For each trial, intracortical local field potential (LFP) signals from −1.0 to 1.5 s around the go cue were converted to μV, downsampled to 500 Hz, and common-median referenced within each 128-channel array. For a given array and trial, CMR was computed only when at least 96 channels were neither constant nor saturated. Beta (13-30 Hz) and high-gamma (70-150 Hz) analytic powers were estimated using zero-phase band-pass filtering followed by Hilbert transformation. Narrow 100- and 150-Hz notches were applied before estimating high-gamma power. Power was expressed in dB relative to the −0.8 to −0.3 s baseline. The primary task response was defined as the mean power change from 0 to 0.8 s after the go cue. Trial-level power curves were first summarized within each physical array and then across the 15 implanted arrays to obtain one session-level curve. Session effects were reported as medians across the 15 implanted arrays with array-level bootstrap 95% confidence intervals. All 180 array-session combinations passed quality control. Of these, 175 retained all 60 selected trials, while five retained 59 trials after artifact exclusion. To quantify changes during chronic implantation, a Theil-Sen slope was estimated separately for each array across the twelve sessions. The median array-level slope and its array-level bootstrap 95% confidence interval were then reported.

Beta-band responses varied across sessions (**Figure 6**A, C). Median changes ranged from −1.03 dB on day 55 (95% CI, −1.21 to −0.69) to 0.24 dB on day 115 (0.11 to 1.57), with no consistent direction across the twelve sessions. Thus, this analysis did not demonstrate reproducible beta event-related desynchronization. The median array-level slope was positive (0.15 dB/30 days; 95% bootstrap CI, 0.12 to 0.30), reflecting a shift toward more positive beta modulation across the selected time points. However, because the session-level responses were non-monotonic and the go cue did not represent movement onset, this slope was not interpreted as a progressive change in movement-related beta suppression.

**Figure 6.**
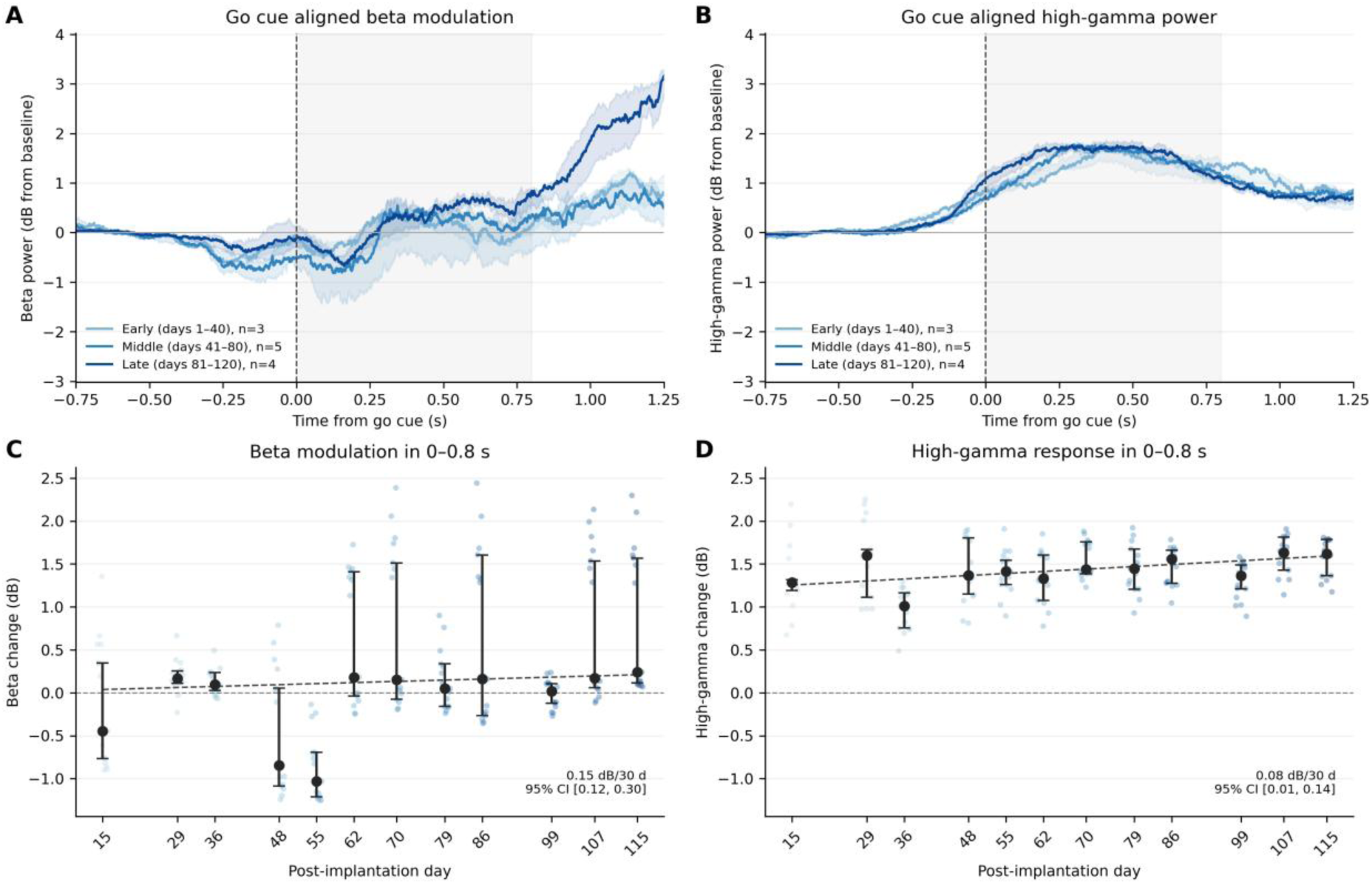
Task-related beta and high-gamma modulation in twelve representative sessions. (A, B) Go-cue-aligned beta and high-gamma power during the early (days 1-40; n=3 sessions), middle (days 41-80; n=5), and late (days 81-120; n=4) periods. Within each session, curves were summarized across the 15 implanted arrays; lines denote the median across sessions in each period and shading denotes the interquartile range across sessions. The grey region marks the prespecified 0-0.8 s response window. (C, D) Array-level beta and high-gamma changes averaged over that response window and plotted against post-implantation day. Colored points denote individual arrays; black points and error bars denote the across-array median and array-level bootstrap 95% confidence interval, and dashed lines denote Theil-Sen trends through the session medians. Power is expressed in dB relative to the −0.8 to −0.3 s baseline. Sixty successful trials were selected per session, and all array-session observations retained at least 59 trials.

High-gamma power increased consistently after the go cue in all twelve sampled sessions (**Figure 6**B, D). The median 0-0.8-s response ranged from 1.01 dB on day 36 (95% CI, 0.75 to 1.16) to 1.63 dB on day 107 (1.43 to 1.81), and every session-level confidence interval was above zero. The median array-level slope across the twelve sampled sessions was 0.08 dB/30 days (95% bootstrap CI, 0.01 to 0.14). The time-locked high-gamma response therefore remained detectable across the sampled chronic implantation period.

### 4. Accuracy of markerless 3D hand-pose reconstruction model

We evaluated the 3D hand-pose reconstruction model by measuring the 3D localization error of 22 keypoints, with Wrist_medial and Wrist_lateral excluded. Model acceptance required a median 3D localization error of no more than 10 mm for the elbow and no more than 5 mm for each of the other 21 keypoints. Following repeated training with successive refinements to the training dataset, the final model met these acceptance criteria on the test set, and visual inspection showed that its predicted poses were generally consistent with the manual annotations. The resulting error array contained 400 frames and 22 keypoints per frame, yielding 8,800 finite frame-keypoint distances with no missing or non-finite values. Because the distributions were right-skewed, performance was summarized using the prespecified keypoint-wise median error rather than the arithmetic mean of all frame-level errors (**Figure 7**A).

**Figure 7.**
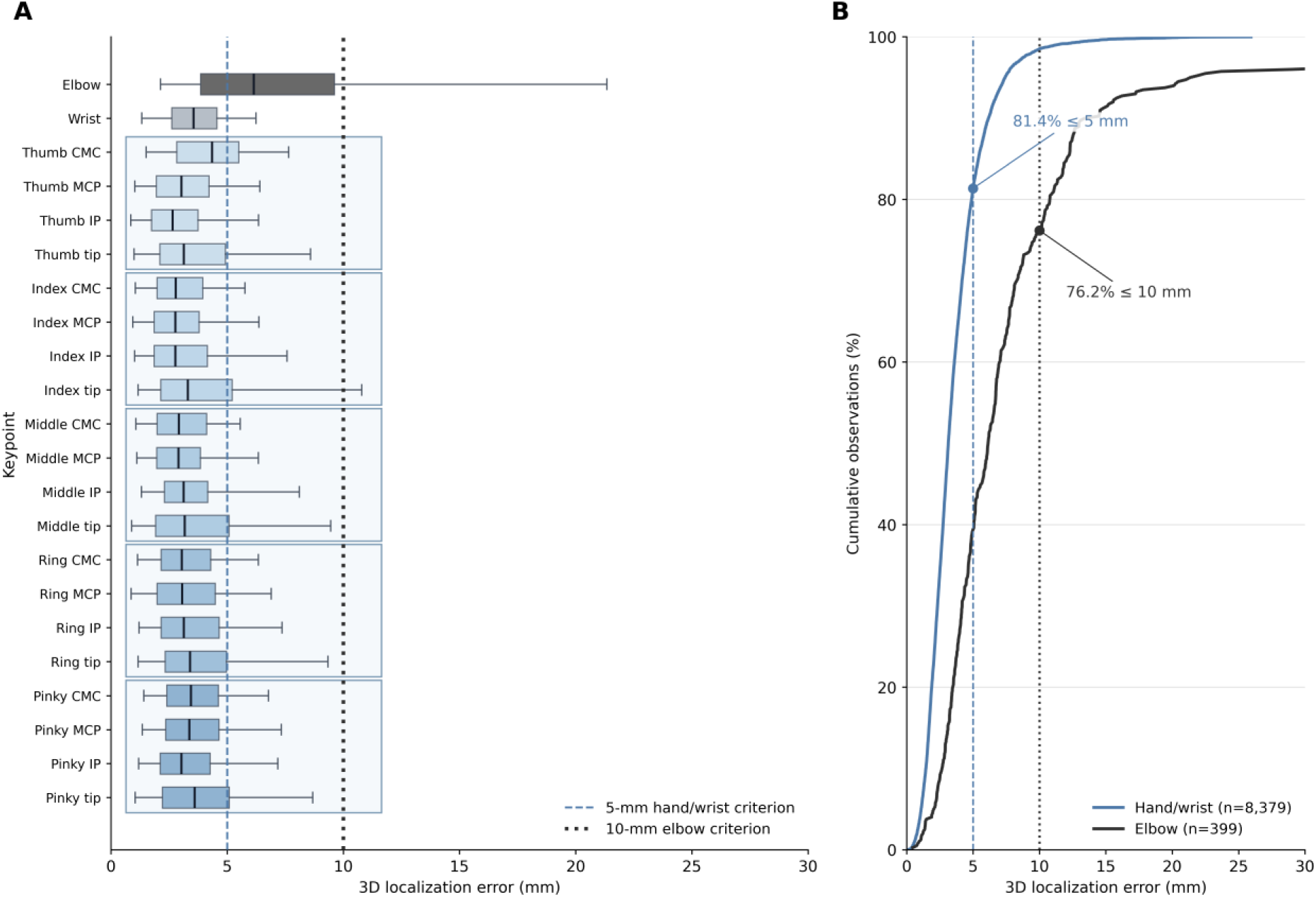
Three-dimensional localization error of the markerless hand-pose model. (A) Frame-level Euclidean errors for each of the 22 keypoints across 400 evaluation frames. Boxes show the median and interquartile range, and whiskers show the 5th and 95th percentiles. Light blue rectangular outlines group the four keypoints of each finger to distinguish the five fingers visually. Observations outside the whiskers were retained in the analysis but are not drawn individually. Dashed segments mark the prespecified 5-mm criterion for the 21 hand and wrist keypoints and the 10-mm criterion for the elbow. (B) Empirical cumulative distributions of the 8,379 pooled hand and wrist distances and 400 elbow distances. Filled circles mark the proportions meeting the corresponding criteria. The displayed x-axis is limited to 30 mm for readability. The maximum observed error was 81.062 mm. Errors were calculated without rigid alignment using the camera calibration shared by the predictions and manual annotations.

All 21 hand and wrist keypoints had median 3D localization errors satisfied the acceptance criterion, ranging from 2.648 mm for the thumb IP to 4.348 mm for the thumb CMC. The elbow median was 6.139 mm. The unweighted mean of the 22 keypoint-specific medians was 3.302 mm. At the individual frame-keypoint level, 81.37% of the 8,379 hand and wrist distances were no greater than 5 mm, and 76.19% of the 400 elbow distances were no greater than 10 mm (**Figure 7**B). Thus, the prespecified criteria were met at the keypoint-median level, although the right-tailed distributions show that some individual frames had substantially larger errors and still require frame-level quality control.

### 5. Neural decoding benchmark

The benchmark compared 12 neural decoders across 20 recording sessions. Within each session, condition-stratified trials were divided into 80% training and 20% testing subsets, and the split was repeated 10 times. This design yielded 200 train-test evaluations per decoder. Repeat-level estimates were averaged within each session before cross-session aggregation. The Hand Reconstruction Fidelity Score (HRFS) together with Macro R², Macro Pearson r, root mean squared error (RMSE), mean absolute error (MAE), and mean per-joint position error (MPJPE) were used as the evaluation metrics.

LMLF achieved the highest cross-session HRFS, with a mean of 58.09 and a bootstrap 95% CI of 56.15 to 59.93. The strongest comparator was long short-term memory (LSTM) regression, which obtained an HRFS of 51.58 (95% CI, 49.08 to 53.94), followed by gated recurrent unit (GRU) regression at 49.91 (95% CI, 47.61 to 52.12) and simple recurrent neural network (RNN) regression at 45.07 (95% CI, 42.47 to 47.54). Thus, LMLF exceeded LSTM by 6.51 HRFS points, corresponding to a relative improvement of 12.63%. Among the remaining methods, selected-LFADS with a linear readout and the Kalman filter achieved mean HRFS values of 35.83 and 34.45, respectively. The classical regression methods ranged from 29.38 for PCA-Ridge regression to 32.57 for support vector regression (SVR).

Decomposition of HRFS further separated performance into global translation, wrist-centered pose, local motion, and skeletal consistency. LMLF achieved component scores of 62.99 for global translation, 62.58 for wrist-centered pose, 33.24 for local motion, and 79.84 for skeletal consistency. LSTM obtained corresponding scores of 56.76, 55.62, 27.84, and 71.93. Therefore, LMLF improved the wrist, pose, motion, and bone components by 6.23, 6.96, 5.39, and 7.91 points, respectively. Skeletal consistency yielded the highest score, whereas local motion remained the lowest-scoring component, identifying temporal movement fidelity as the principal remaining source of error.

The session-level comparison in **Figure 8**A shows that LMLF ranked first in all 20 recording sessions. Relative to LSTM, the closest competing method, the session-wise advantage ranged from 3.81 to 10.25 HRFS points and averaged 6.51 points. Although the absolute scores varied between sessions, LMLF preserved its leading position throughout the complete session set.

**Figure 8.**
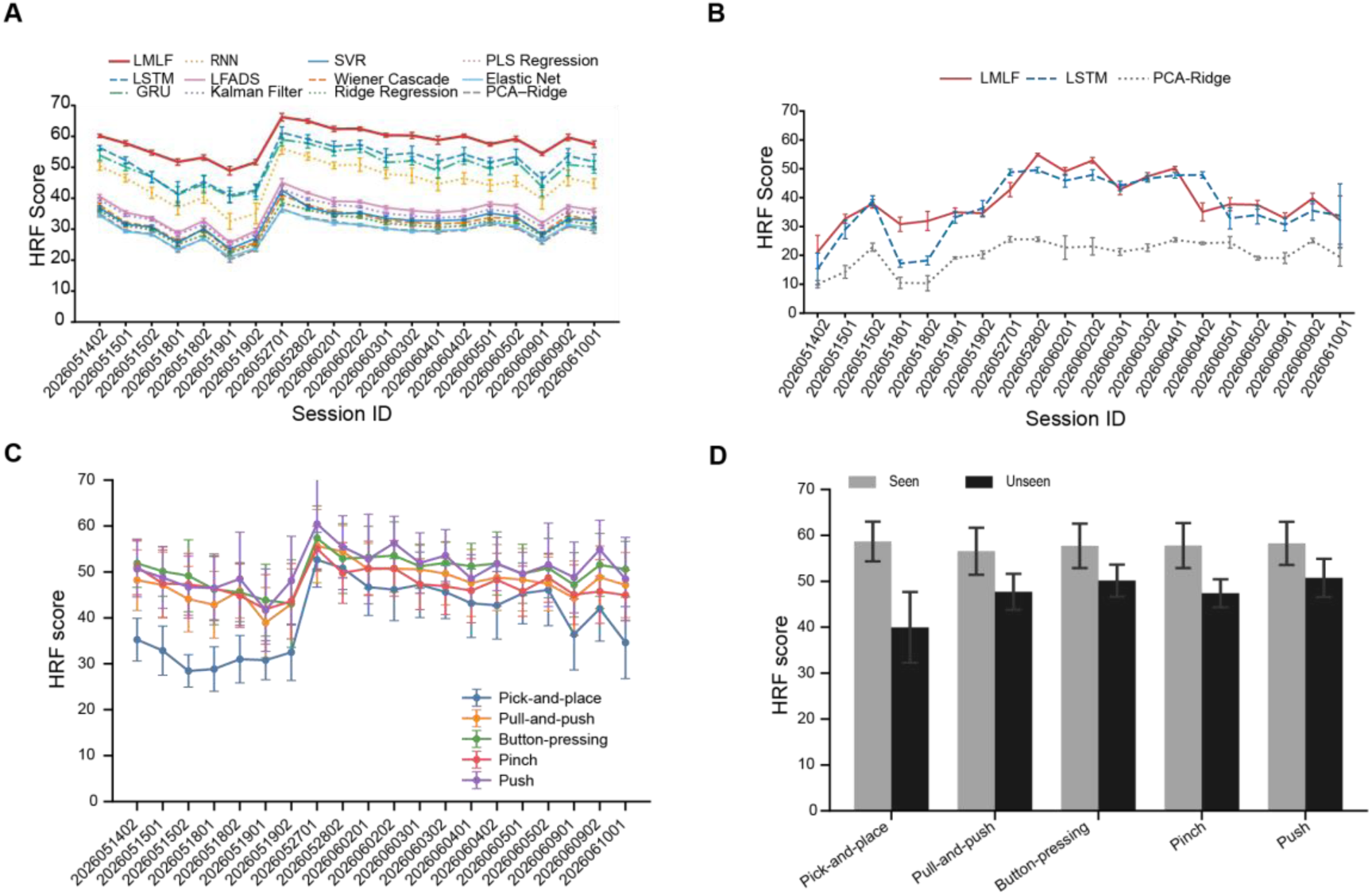
Neural decoding performance and generalization test. (A) Session-level HRFS for the 12 benchmark decoders. Curves show the mean across 10 repeated holdouts. Error bars show the corresponding standard deviation. LMLF is highlighted in red. (B) Cross-day hand-keypoint decoding performance of LMLF, LSTM, and PCA-Ridge under leave-one-recording-day-out evaluation. All sessions from the test day were excluded from training. Curves show the mean HRFS for each held-out session. Error bars indicate ±1 standard deviation across ten repeated source-training subsamples. Test trials were fixed across repeats, so that the error bars reflect variability caused by the influence of training-sample selection on decoders. (C) Cross-task hand-keypoint decoding performance of LMLF across sessions and held-out task categories. Curves show the mean across 10 repeated experiments, and error bars indicate the corresponding standard deviation. (D) Comparison of cross-task hand-keypoint decoding performance of LMLF between seen and unseen test sets for each held-out task category. Bars show the mean across sessions, and error bars indicate the standard deviation across sessions.

The conventional metrics yielded the same overall conclusion as HRFS (Table 5). LMLF achieved a Macro R² of 0.905 and a Macro Pearson r of 0.952, compared with 0.863 and 0.930 for LSTM. LMLF also produced the lowest reconstruction errors, with an RMSE of 10.73, an MAE of 6.26, and an MPJPE of 13.00. The corresponding LSTM values were 12.84, 7.66, and 15.81. Relative to LSTM, LMLF reduced RMSE by 16.49%, MAE by 18.29%, and MPJPE by 17.81%. These concordant improvements across scale-dependent errors, explained variance, and temporal correlation indicate that the HRFS gain was accompanied by broader improvements in coordinate-level reconstruction accuracy.

**Table 5.** Aggregate decoding performance across 20 recording sessions.

| Method | HRFS | Macro $R^2$ | Macro Pearson $r$ | RMSE (mm) | MAE (mm) | MPJPE (mm) |
| --- | --- | --- | --- | --- | --- | --- |
| **LMLF** | **58.09** | **0.905** | **0.952** | **10.73** | **6.26** | **13.00** |
| LSTM | 51.58 | 0.863 | 0.930 | 12.84 | 7.66 | 15.81 |
| GRU | 49.91 | 0.855 | 0.926 | 13.22 | 8.14 | 16.78 |
| Simple RNN | 45.07 | 0.822 | 0.907 | 14.67 | 8.94 | 18.44 |
| Selected LFADS + Linear | 35.83 | 0.630 | 0.790 | 21.28 | 14.34 | 29.32 |
| Kalman Filter | 34.45 | 0.624 | 0.799 | 21.41 | 14.61 | 29.76 |
| SVR | 32.57 | 0.547 | 0.736 | 23.63 | 15.05 | 31.11 |
| Wiener Cascade | 32.34 | 0.272 | 0.770 | 23.58 | 14.74 | 30.39 |
| Ridge | 31.18 | 0.586 | 0.763 | 22.51 | 15.82 | 32.15 |
| PLS | 31.05 | 0.585 | 0.762 | 22.55 | 15.84 | 32.20 |
| Elastic Net | 29.56 | 0.521 | 0.717 | 24.30 | 16.86 | 34.32 |
| PCA-Ridge | 29.38 | 0.105 | 0.704 | 27.40 | 17.13 | 34.79 |

**Table 6.** Results of Cross-task Generalization Test.

| Held-out task category | Seen HRFS | Unseen HRFS | Relative gap (%) |
| --- | --- | --- | --- |
| Pull-and-push | 56.55 ± 5.12 | 47.71 ± 3.92 | 15.53 ± 2.48% |
| Push | 58.26 ± 4.69 | 50.75 ± 4.15 | 12.84 ± 3.20% |
| Pick-and-place | 58.66 ± 4.32 | 39.96 ± 7.70 | 32.27 ± 9.59% |
| Button-pressing | 57.73 ± 4.84 | 50.15 ± 3.49 | 13.00 ± 2.37% |
| Pinch | 57.79 ± 4.90 | 47.39 ± 3.04 | 17.78 ± 3.99% |

LMLF’s higher overall cross-day performance and consistent advantage over PCA-Ridge suggest that it may capture nonlinear relationships between neural activity and hand movement that the linear baseline does not adequately represent. Some of these relationships may be shared across recording days, supporting decoding on unseen days. Nevertheless, the reduction from within-session performance suggests that the learned mapping remains sensitive to day-specific variation, alongside differences in training and evaluation conditions. These findings support assessing decoders not only by within-session reconstruction accuracy but also by their ability to generalize across recording days.

Cross-task generalization also differed substantially across task categories. Pick-and-place had the lowest unseen HRFS, at 39.96, compared with a seen HRFS of 58.66, corresponding to an approximately 31.9% relative performance reduction. This reduction was larger than that observed for the other task categories. In comparison, the relative generalization gaps for pull-and-push, button-pressing, pinch, and push were approximately 12.9%-18.0%, indicating stronger transferability across these categories.

The lower cross-task performance of pick-and-place was observed across multiple recording sessions rather than being driven by a single session (**Figure 8**C). Thus, within the current LMLF framework, the model learned partially shared neural activity-hand movement relationships across task categories, but these shared representations remained constrained by task-specific movement structures. Overall, LMLF showed uneven generalization across task categories, with pick-and-place being the most difficult category to predict from the other tasks. The results of cross-day and cross-task generalization tests emphasize the value of data sets that cover prolonged sampling periods and multiple motor task categories. Such data sets may provide large feature spaces of both movement and neural activity, which is necessary for developing algorithms with robust generalization ability.

### 6. Robustness to reduced channel availability

Under limited channel availability, LMLF had the most favorable mean value among the three decoders at every channel count for all six metrics (Figure 9). With all 1,920 channels, LMLF achieved an HRFS of 58.14, a Macro R² of 0.907, and a Macro Pearson r of 0.953, with RMSE, MAE, and MPJPE values of 10.56 mm, 6.20 mm, and 12.83 mm, respectively. The corresponding full-channel values were 51.37, 0.862, 0.930, 12.91 mm, 7.76 mm, and 16.02 mm for LSTM and 29.42, 0.336, 0.701, 26.44 mm, 16.85 mm, and 34.23 mm for PCA-Ridge.

**Figure 9.**
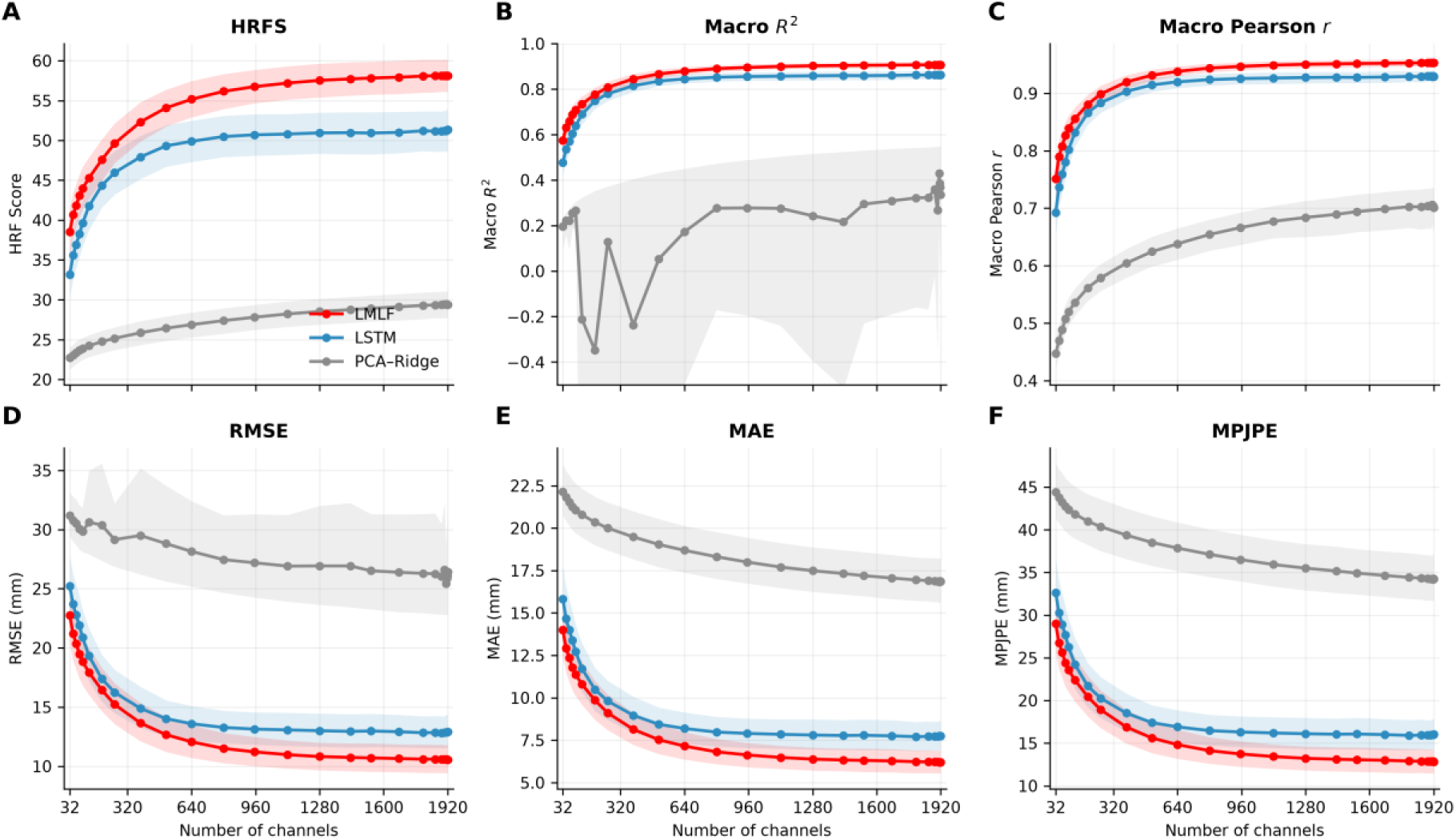
Decoding performance as a function of channel availability. (A) HRFS, (B) Macro R², (C) Macro Pearson r, (D) RMSE, (E) MAE, and (F) MPJPE for LMLF, LSTM, and PCA-Ridge regression. LMLF, LSTM, and PCA-Ridge are shown in red, blue, and gray, respectively. The horizontal axis gives the number of eligible physical channels retained for decoding. Each channel contributed one TX and one SBP feature. The dislodged eg1-8 array was excluded in all conditions. Points show the mean across 20 recording sessions and 50 random channel masks at each non-baseline channel count. Shaded bands show 95% confidence intervals from a two-way bootstrap that independently resampled sessions and masks. The 1,920-channel baseline contained one unmasked configuration. We set the display range of Macro R² as −0.5 to 1.0 to show the mean trends. We clipped the portions of the PCA-Ridge confidence interval that were lower than −0.5. Higher values indicate better performance for HRFS, Macro R², and Macro Pearson r. Lower values indicate better performance for RMSE, MAE, and MPJPE. Error metrics are reported in millimeters.

When 954 channels remained, LMLF retained an HRFS of 56.75, a Macro R² of 0.895, and a Macro Pearson r of 0.947, with RMSE, MAE, and MPJPE values of 11.22 mm, 6.63 mm, and 13.73 mm. The corresponding values were 50.70, 0.855, 0.926, 13.14 mm, 7.90 mm, and 16.30 mm for LSTM and 27.82, 0.277, 0.666, 27.19 mm, 18.00 mm, and 36.50 mm for PCA-Ridge. Thus, LMLF and LSTM both remained close to their full-channel performance when approximately half of the eligible channels were retained, with LMLF maintaining a higher mean performance.

At the most restrictive 32-channel condition, LMLF obtained an HRFS of 38.50, a Macro R² of 0.574, and a Macro Pearson r of 0.751, with RMSE, MAE, and MPJPE values of 22.77 mm, 13.99 mm, and 28.98 mm. LSTM obtained corresponding values of 33.16, 0.477, 0.693, 25.22 mm, 15.83 mm, and 32.63 mm, while PCA-Ridge obtained 22.71, 0.196, 0.447, 31.18 mm, 22.14 mm, and 44.39 mm.

The Macro R² of PCA-Ridge was non-monotonic, and its mean fell below zero at several intermediate channel counts, indicating that squared prediction error exceeded target variance on average in those conditions. Macro Pearson r, HRFS, and the three error metrics showed a clearer loss of performance with decreasing channel availability. These results show that useful kinematic information remained available under substantial channel loss while identifying the range in which limited neural coverage became a dominant constraint. During the design of BCI systems, similar analysis with a high-throughput data set like the current one may help to optimize the hardware and algorithms.

## Data availability

The current release of the dataset with selected trials is available at https://dbci.lglab.ac.cn/competition/dataset/X54wEnWr

## Code availability

The code used for the processing of data and our preliminary analyses is available from the corresponding author, upon reasonable request.

## Acknowledgements

We thank Li Li, Qing Xu and Yuheng Zhang for their support in animal experiments. We thank Tikui Zhang and other staff at the animal facility for expert monkey care. We thank Xinyu Jian, Shi Wang, and Xiaoyang Liu for administrative support. We thank Zhenliang He for his input during discussions of research ideas. We appreciate Yifan Huang for his effort in developing a motion capture system. It inspired our current design of the motion capture system and cross-system synchronization. Nicolas Zdun at the German Primate Center kindly provided useful suggestions related to the use of JARVIS-Mocap. This work was supported by Lingang Laboratory (Grant No. LGL-7209, Grant No. LG-GG-202402-06).

## Author information

Tianmeng He designed the behavioral paradigm with guidance from Chengyu T. Li, Yang Xie and Yanan Sui. Tianmeng He, Kai Liu and Shuran Guan designed and built the experimental setup as well as the data recording systems. Shouliang Guan developed the flexible microelectrode arrays. Yulei Chen, Hongxiong Jiang, Lei Wu, Ruoyu Li and Fengmei Zhu conducted surgeries on the subject. Mengmeng Chen, Dongling Zhang, Long Chen and Shundong Huang trained the monkeys to perform the task, and conducted electrophysiological data and behavioral data recordings. Sizheng Luo, Yahui Li, Yizhi Dong, Weitao Chen, Kai Liu and Shuran Guan set up the data processing workflow and performed analysis with guidance from Tianmeng He, Tianwei Wang, Yanan Sui, Bin Min, and Chengyu T. Li. Sizheng Luo, Kai Liu, Shuran Guan and Tianmeng He created the figures. Sizheng Luo, Yahui Li, Kai Liu and Shuran Guan wrote the initial manuscript and contributed to editing as well as revision. Tianmeng He and Tianwei Wang contributed to editing and revisions of the manuscript. Tianmeng He, Tianwei Wang and Chengyu T. Li jointly supervised and coordinated the project.

